# Morphological and phylogenetic resolution of *Conoideocrella luteorostrata* (Hypocreales: Clavicipitaceae), a potential biocontrol fungus for *Fiorinia externa* in United States Christmas tree production areas

**DOI:** 10.1101/2022.10.18.512709

**Authors:** Brian Lovett, Hana Barrett, Angie M. Macias, Jason E. Stajich, Lindsay R. Kasson, Matt T. Kasson

## Abstract

The entomopathogenic fungus *Conoideocrella luteorostrata* (Hypocreales: Clavicipitaceae) has recently been implicated in natural epizootics among exotic elongate hemlock scale (EHS) insects in Fraser fir Christmas tree farms in the eastern U.S. Since 1913, asexual populations of *C. luteorostrata* have been reported from various plant-feeding Hemiptera in the southeastern U.S., but a thorough morphological and phylogenetic examination of the species, particularly detailed characterization of populations involved in recent epizootics in EHS, are lacking. The recovery of multiple strains of *C. luteorostrata* from mycosed EHS cadavers collected in Ashe County North Carolina provided an opportunity to conduct pathogenicity assays and morphological and phylogenetic studies to investigate genus- and species-level boundaries among members of the Clavicipitaceae. Pathogenicity assays confirmed *C. luteorostrata* causes mortality of EHS first instar crawlers, an essential first step in developing *C. luteorostrata* as a biocontrol. The results of the morphological study failed to recover a sexual stage from EHS cadavers or pure cultures, but revealed conidia aligned with previous measurements of the paecilomyces-like asexual state of *C. luteorostrata* (6.9 µm x 2.6 µm average), with colony and conidiophore morphology consistent with previously reported observations. Additionally, a hirsutella-like synanamorph of *C. luteorostrata* was observed for the first time under specific lab conditions. In both a four-locus, 54-taxa Clavicipitaceae-wide phylogenetic analysis including the D1–D2 domains of the nuclear 28S rRNA gene (28S), elongation factor 1 alpha (EF1-α), DNA-directed RNA polymerase II subunit 1 (RPB1) and DNA-directed RNA polymerase II subunit 2 (RPB2) and a two-locus, 38-taxa (28S & EF1-α) phylogenetic analysis, *C. luteorostrata*, *C. tenuis* and *C. krungchingensis* were resolved as strongly supported monophyletic lineages across all loci and both methods (maximum likelihood & Bayesian inference) of phylogenetic inference with the exception of 28S for *C. tenuis*. Despite the strong support for individual *Conoideocrella* species, none of the analyses supported the monophyly of the genus, with the inclusion of *Dussiella*. Due to the paucity of publicly available RPB1 and RPB2 sequence data for *Conoideocrella*, EF1-α provided superior delimitation of intraspecies groupings for *C. luteorostrata* and *C. tenuis* and should be used in future studies. Further development of *C. luteorostrata* as a biocontrol agent against EHS both in Christmas tree farms and surrounding hemlock forests will require additional surveys across diverse Hemiptera and expanded pathogenicity testing to better understand host range and efficacy of this fungus.

## Introduction

In the summer of 2020, the entomopathogenic fungus *Conoideocrella luteorostrata* (CL) was found causing an emergent epizootic on elongate hemlock scale insects (EHS; *Fiorinia externa*) infesting Fraser fir (*Abies fraseri*) trees on Christmas tree farms in North Carolina. EHS was first documented in the U.S. in 1908 on eastern hemlock (*Tsuga canadensis*) in New York State (Sasscer, 1912), but this non-native pest has since spread throughout the eastern U.S. The range of EHS was originally limited by their cold sensitivity, but adaptations that increased cold tolerance emerged in the 1970s which allowed for rapid northward expansion (Preisser et al., 2008).

Despite its name, confirmed plant hosts of EHS include spruce, fir, and pine species, in addition to eastern hemlock (McClure & Fergione, 1977). Despite its long residency on native hemlock, EHS was first reported in Christmas tree production areas on Yancey County North Carolina Fraser fir tree farms around 1993 (Miller & Davidson 2005; Sidebottom, 2016). The damage inflicted by these insect pests on true fir tree farms and on eastern hemlock in native forested stands is challenging to quantify, but the spread of this insect, along with another non-native insect pest hemlock woolly adelgid (HWA; *Adelges tsugae*), has been associated with defoliation and decline of hemlocks in the United States (Royle & Lathrop, 2002). Given the broad host range of EHS, this pest poses significant risks to the Christmas tree production industry, where scale infection can decrease the ability of farmers to transport and sell their products across state lines (McClure & Fergione, 1977), since the transport of Christmas trees can contribute to the spread of this and other non-native insect pests (Dale et al., 2020).

Chemical control has limited efficacy against EHS and other armored scales. Due to the high fertility of EHS and the vulnerability of their natural predators to chemical treatments, the pest population can rebound to higher levels after treatment (McClure, 1977a). Fungal biocontrol agents have several characteristics that could be especially valuable for the management of EHS. Since entomopathogenic fungi produce diverse enzymes, toxins and other secondary metabolites targeting their hosts as well as diverse mechanisms for overcoming host immune defenses, scales are unlikely to easily evolve resistance to fungal pathogens (Gao et al. 2017). Fungal biocontrol agents are often highly host-specific, so they may have fewer non-target effects than chemical pesticides (McClure, 1977b). The fungus *Colletotrichum fioriniae* has been proposed as a biological control agent of EHS on eastern hemlock (Marcelino et al., 2009), but further research showed that it is primarily a leaf/stem endophyte (Martin & Peter, 2021). *C. fioriniae* also has been reported as a post-harvest pathogen of apples (Chechi et al., 2019), other fruits (Ling et al., 2021; Ivic et al., 2013), and native plants (Kasson et al., 2014), potentially limiting its potential as a biocontrol agent for EHS. Another putative fungal pathogen of elongate hemlock scale, *Metarhiziopsis microspora*, has also been described from the United States by Li and colleagues (2008), but its pathogenicity, ecology, and non-target effects have not been further characterized. If a fungal biocontrol agent can be developed for EHS, it would be a valuable addition to an integrated pest management plan for Christmas tree growers. This biopesticide would also support resource managers of natural forested areas where eastern hemlock trees are succumbing to the cumulative effects of HWA and EHS.

In addition to confirmed epizootics in infested Fraser fir Christmas trees in North Carolina, CL also has been recently reported from mycosed EHS on these same hosts in Virginia and Michigan (Urbina & Ahmed, 2022). Formerly known as *Torrubiella luteorostrata* (teleomorph or sexual form) or *Paecilomyces cinnamomeus* (PC; anamorph or asexual form), this fungus was first observed in the U.S. on whiteflies in Louisiana in 1913, followed by Mississippi in 1920, Alabama in 1923 and Florida in 1937 (TABLE 1; Samson, 1974). Outside the U.S., both the asexual and sexual states of this fungus have been reported from several scale and whitefly species spanning three families in the hemipteran suborder Sternorrhyncha, from a total of thirteen countries (including the US) distributed across Africa (Ghana, Kenya and Seychelles), Asia (China, Indonesia, Japan, Sri Lanka and Thailand), North America (Cuba, Mexico and the U.S.), Oceania (Samoa) and South America (Guatemala) (TABLE 1). Reports of CL in Russia are also known from numerous hosts, yet no morphological or DNA sequence data are available to resolve those populations among known *Conoideocrella* species. Herbarium specimens for CL also exist in public collections (SUPPLEMENTARY TABLE 1) for additional countries including Fiji, New Zealand and Portugal, but no morphological data or DNA sequence data is available, nor were these specimens used in any taxonomic studies on the genus to confirm tentative identifications (TABLE 1). Nevertheless, these publicly archived herbarium specimens represent the only records for CL on members of the families Coccidae and Aleyrodidae in the U.S. and push back the first occurrence of this fungus in the U.S. by more than a century (TABLE 1). One of the commonly reported insect hosts for PC in the early 20th century, *Dialeurodes citri* or citrus whitefly, was itself introduced into Florida from southeast Asia sometime between 1858 and 1885 (Morrill & Back 1911) and may have harbored CL upon arrival.

The genus *Conoideocrella* contains three formally described species: *C. tenuis* (Petch) D. Johnson, G.H. Sung, Hywel-Jones & Spatafora 2009 (CT), *C. krungchingensis* Mongkols., Thanakitp. & Luangsa-ard, 2016 (CK) and *C. luteorostrata* (Schoch et al., 2020). CL is commonly found infecting multiple pest species in the hemipteran suborder Sternorrhyncha, including armored scale insects (Diaspididae), soft scales (Coccidae) and whiteflies (Aleyrodidae), while CK has only been reported from Diaspididae from Thailand. CT infects hosts from Aleyrodidae and Diaspididae, but across a limited geographic area mostly confined to southeast Asia (TABLE 1). CL forms characteristic ochre-to-cinnamon colored stromata upon infected insect hosts with hypothalli consisting of thin hyphae expanding outward radially upon which conidiophores and/or perithecia are formed. CT and CK also produce variously colored stromata and hypothalli, but also have features that are distinct from all previously published accounts of CL, including a hirsutella-like asexual stage in CK with 3-4 transversely septate conidia produced singly and no anamorph observed for CT (Johnson et al., 2009; Saito et al., 2012; Mongkolsamrit et al., 2016).

*Dussiella tuberiformis* (Berk. & Ravenel) Pat. ex Sacc. (previously known as *Echinodothis tuberiformis* and *Hypocrea tuberiformis*)—a closely allied member of the Clavicipitaceae (Kepler et al., 2012)—also infects scale insects on switchcane (*Arundinaria tecta*) and has been historically reported from the southeastern U.S. including Alabama, Mississippi, North Carolina and South Carolina (TABLE 1; Atkinson, 1891; Korosh et al., 2004; White et al., 2002). Contemporary observations of *D. tuberiformis* exist on the community science platform iNaturalist.org, and despite some variability in their macroscopic features, are markedly distinct from *Conoideocrella* based on the formation of stroma with crowded aggregations of individual perithecia (Atkinson, 1890; SUPPLEMENTARY TABLE 2). Kepler and colleagues (2012) resolved *Dussiella* as sister to *Conoideocrella*. However, the phylogenetic position of *Dussiella* among *Conoideocrella* species and other members of the Clavicipitaceae remains unclear, given its absence from the recent taxonomic description of CK (Mongkolsamrit et al., 2016). Two other species of *Dussiella, D. orchideacearum* Rick 1906 and *D. violacea* Höhn 1907, have been previously described, but neither has publicly available sequence data to resolve their relationships among *Conoideocrella* and *D. tuberiformis* (TABLE 1).

Given the taxonomic uncertainty between *Conoideocrella and Dussiella*, and the potential of developing a U.S. strain of CL as a biocontrol agent against EHS and potentially against non-native whiteflies, this study was undertaken to: 1) provide a comprehensive summary of previous work on these fungi; 2) morphologically and phylogenetically resolve the recently discovered U.S. populations of CL among previously characterized *Conoideocrella* and *Dussiella*; and 3) conduct pathogenicity testing on EHS to further explore its development as a potential biocontrol agent against this invasive insect. Though numerous surveys around the globe have confirmed the presence of CL, CT and CK on various hemipteran hosts, robust multi-locus phylogenetic analyses coupled with formal investigation of pathogenicity of CL on armored scale insects has not been previously pursued.

## Methods

### Surveying for fungal pathogens of elongate hemlock scale

Fungal surveys targeting entomopathogenic fungi on elongate hemlock scale (EHS) were conducted at three sites in Ashe County, North Carolina (Vannoy Farm (VF), Upper Mountain (UM), and Deal Family Farm (BAF)) and two sites in Grayson County, Virginia (Mount Rogers (MR) and Mount Rogers Orchard (MRO)) in late August and early September, 2020. Though VF and BAF are on private lands with limited access, a third site, UM, serves as a research station for North Carolina State University’s College of Agriculture and Life Sciences and is publicly accessible for research. Coordinates associated with each sampling location are provided in SUPPLEMENTARY TABLE 3. All three of the North Carolina sampling locations were reported as moderate to heavy EHS infestations and none were actively managed with chemical controls. EHS infestations had not been reported from either site at Mount Rogers prior to this survey. Infected EHS were identifiable by the distinct orange mat of hyphae covering the cadavers, and were abundant, with multiple infected cadavers on each infested needle (FIG. 1A-B). EHS-free branches were collected from all sites, as were branches with uninfected insects from the four EHS positive sites, and branches with mycosed insects from the three North Carolina sites. Samples were individually bagged, such that insects found among branches were also preserved by site to allow for subsequent examination.

**Figure 1.**
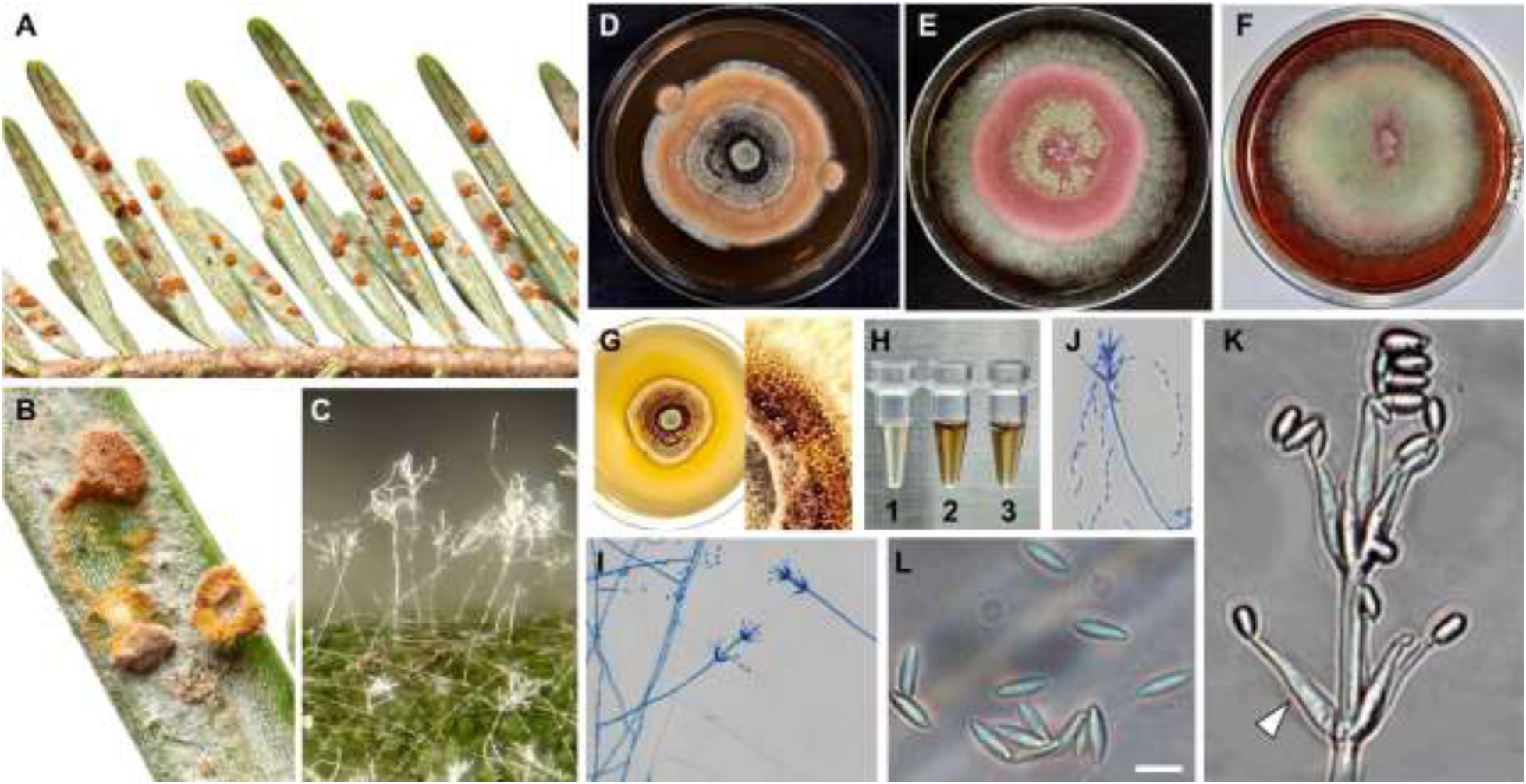
Natural infections (A-C), pure cultures (D-G) and select morphological features (I-L) including extracted pigments (H) from *C. luteorostrata* strains recovered from an epizootic on EHS infesting Fraser fir Christmas trees in North Carolina. Macroscopic features include orange stroma produced over killed elongate hemlock scale crawler cadavers on the underside of fir needles (A-B) and and verticilliate condiophores (C) emerging from the white mycelium immediately surrounding the mycosed EHS. Representative cultures on PDA+ST at 1-2 weeks (D), 3-4 weeks (E), and 5+ weeks of age (F). Pigment production by CL on PDA (G) and following extraction from 1.78 cm diameter colonized agar plugs from a >5+ week old culture with water (H1), ethanol (H2), and methanol (H3). Microscopic features include representative conidiophores (C, I-K) with phialides (white arrow), K) and conidia (J-L) mounted in lactic acid with and without cotton blue. Scale bars = 5 µm for panel L. Photos in panel A-C are from Watauga County, North Carolina and taken by Dr. Matt Bertone.

**Figure 2.**
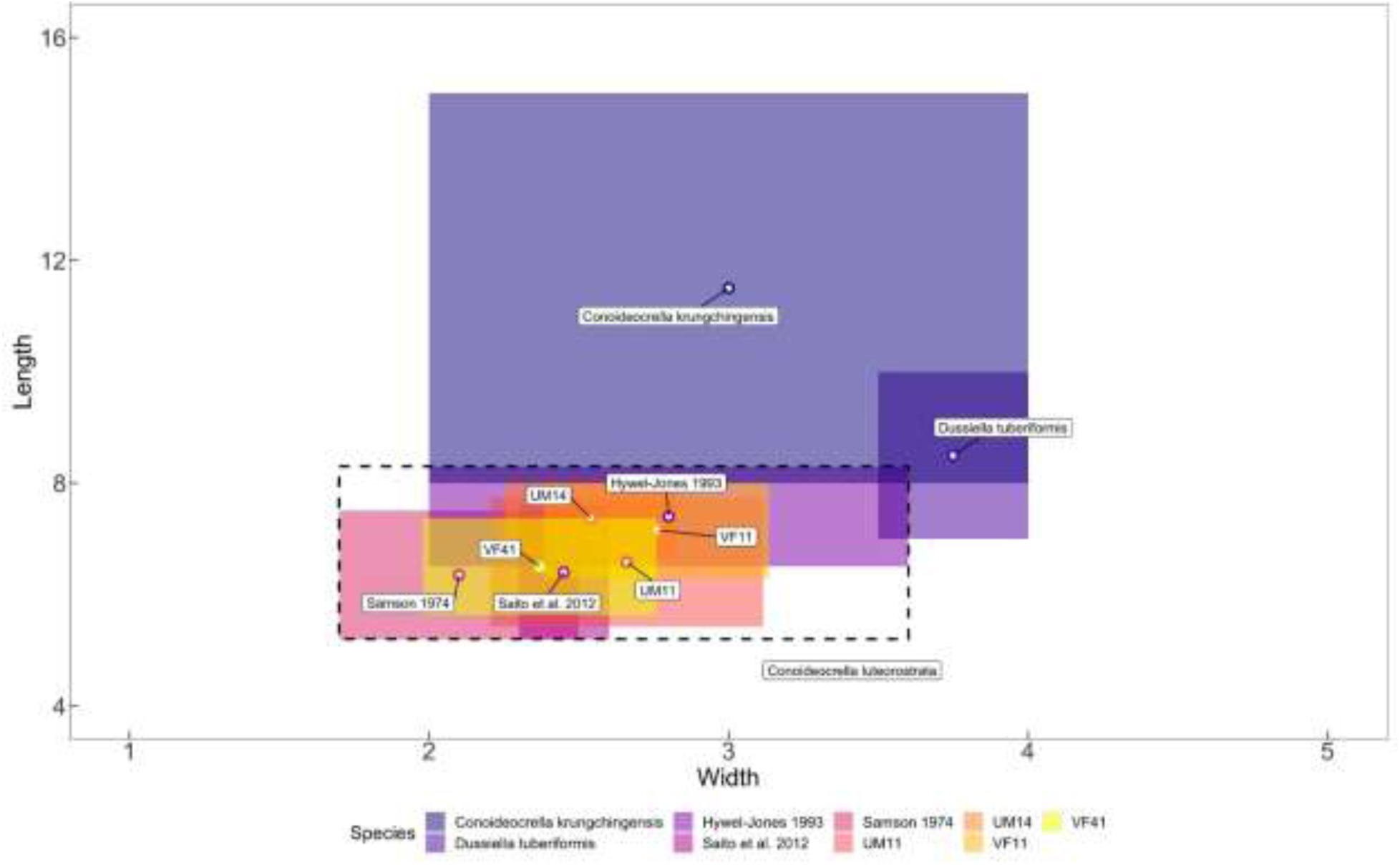
Primary conidial length and width measurements (µm) for strains from North Carolina overlaid atop measurements previously reported for *C. luteorostrata* (inside black dashed line), *C. krungchingensis*, and *D. tuberiformis*. Mean values are denoted by a circle. For CL measurements, names following colored box denotes paper from which the measurements were pulled. All other CL strain data was generated as part of this study. Measurements also available in Table 4 and Supplementary Table 5.

### Fungal isolations

Fungi were cultured from EHS by removing the infected/mycosed nymph cadavers from fir needles with a sterile scalpel or teasing needle and placing them on a potato dextrose agar (PDA; Difco, Detroit, Michigan) plate supplemented with 0.1 g/L streptomycin and 0.01 g/L tetracycline (+ST) to suppress bacterial growth. Cultures were prepared from representative samples from each location with infected EHS. Single colonies emerging from plated infected nymphs up to 2 weeks post-plating were subcultured and retained in pure culture until colonies were of sufficient size to compare morphotypes. A single dominant morphotype (∼85% of all colonies) with morphological characteristics consistent with the paecilomyces-like asexual state of CL, including ochre-to-cinnamon colored mycelia with verticillate conidiophores and the production of a reddish pigment secreted into the agar by the fungus (Saito et al., 2012), was observed and retained for further morphological and molecular investigation (FIG. 1).

### Culture-based survey of fir needles

Because other hemipteran entomopathogens, such as *Colletotrichum fioriniae* (EHS) and *Dussiella tuberiformis*, are known leaf endophytes exhibiting epibiont growth on their plant hosts (Martin & Peter, 2021; Korosh et al., 2004; White et al., 2002), we sought to determine if CL could be re-isolated from healthy and dead Fraser fir needles sampled from trees with and without EHS and/or CL. The culturable fungal communities of the fir needles were studied from trees and needles with no EHS, EHS only, and EHS infected with CL. A total of sixty needles were sampled across 5 previously referenced sampling sites. This included 24 live needle samples from six EHS+CL+ trees (UM & VF), 12 samples from three EHS+CL-trees (MRO), 12 samples from three EHS-CL-trees (BAF) and 12 dead needle samples from one EHS-CL-tree (MR) and two trees at BAF.

Needles were sterilized in 10% sodium hypochlorite for 20 seconds followed by a sterile water rinse for 20 seconds, placed on sterile filter paper to dry for one minute, and then plated onto PDA+ST. Isolated fungi were retained in pure culture to compare morphotypes, and representatives of the most prevalent morphotypes along with any morphotypes that had morphological characteristics similar to CL were retained for sequencing.

### Superficial survey for fungal infections in co-occurring arthropods

To assess for potential non-target (non-EHS) impacts, living and deceased non-sessile arthropods (i.e., insects, arachnids, and collembolans) found in EHS-infested and EHS-free Fraser fir branch materials were collected, examined under a dissecting scope for conspicuous fungal infections, and once determined to not have obvious outward fungal growth, stored in ethanol for subsequent morphological identification. No fungal isolations were attempted from outwardly asymptomatic living arthropods collected during this survey, nor were collected dead arthropods, which lacked outward infections, incubated for fungal cultivation since CL produces conspicuous, dense growth on infected cadavers. A higher-level identification (i.e., class, subclass, or order) was assigned to all arthropods and, where possible, a lower-level (down to species) identification was provided leveraging resources such as BugGuide and iNaturalist (SUPPLEMENTARY TABLE 4).

### Morphological studies

A total of 13 CL strains recovered from mycosed EHS, spanning three locations in North Carolina, were assessed. Morphological studies included sporulation observations, conidial measurements, and colony growth rates. One representative strain, CLUM14, was deposited in the ARS Collection of Entomopathogenic Fungal Cultures (ARSEF) and will be referred to hereafter as ARSEF 14590.

Slides for measuring spores were prepared from two-week-old PDA+ST cultures of four CL strains: ARSEF 14590, UM11, VF11, and VF14. Slides were prepared with lactic acid as a mountant, sealed with clear nail polish and stored flat for several weeks before measurements were conducted. Slides were examined and photographed using a Nikon Eclipse E600 compound microscope (Nikon Instruments, Melville, New York) equipped with a Nikon Digital Sight DS-Ri1 high-resolution microscope camera. Lengths and widths for 25 conidia for each strain were measured using Nikon NIS-Elements BR3.2 imaging software. Raw spore measurements are available in SUPPLEMENTARY TABLE 5.

Spore measurement data were analyzed using the packages tidyverse (v. 1.3.2; Wickham et al., 2019), agricolae (v. 1.3.5; de Mendiburu, 2021), RColorBrewer (v. 1.1.3; Neuwirth, 2022), scales (v. 1.2.1; Wickham & Seidel, 2022), car (v. 3.1.2; Fox & Weisberg, 2019) and viridis (v. 0.6.2; Garnier et al., 2021) in R (v. 4.2.2; R Core Team, 2022). Normality was assessed using the Shapiro-Wilk test and equality of variance was assessed using Levene’s test. Analysis of variance (ANOVAs) were performed to check for differences in spore measurements across strains, and Tukey’s honest significance test was used to identify any significant pairwise differences. A p-value <0.05 was considered significant for all analyses. All data and code for analyses included in this manuscript are available on GitHub: https://github.com/HanaBarrett/EHS-CL-Analysis.

Colony growth rate was measured to determine temperature optima for CL strains from North Carolina. A sterile 1.78 cm diameter cork-borer was used to cut colonized agar plugs from two-week-old fungal cultures of two strains: BAF14 and VF45. These agar plugs were plated in the center of 9-cm diameter Petri plates containing PDA+ST. Five plates each for each of the two strains were placed in incubation chambers set to 10°C, 22°C, or 30°C and kept in complete darkness. The perimeter of the colony from the center of the petri plate was marked along 4 pre-drawn radii (90°, 180°, 270°, 360°) every two days. After two weeks, the distance between each point and the perimeter of the inoculation plug was measured and recorded. Average daily growth rate was calculated as half the distance of mean 2-day growth rate, the latter of which was the average of individual measurements from 4 radii (i.e., during days 4, 6, 8, 10, 12 and 14). Plates with satellite colonies or atypical radial growth were excluded. For isolates that showed no radial growth between successive time points at a given tested temperature, plates were still monitored for 14 days, after which plates were returned to 22°C to assess whether growth resumed. Raw colony growth measurements are available in SUPPLEMENTARY TABLE 6.

Alternative culturing methods to stimulate development of other additional spore types/asexual states (e.g. hirsutella-like septate conidia reported in CK) and specialized fungal structures (e.g. appressoria) were initiated by observing growth upon a coverslip. This was initiated by two methods. ARSEF 14590 conidia were mixed into a cooling solution of PDA, after which a small volume (∼5 µm) was placed on a sterile slide beneath a sterile cover slip. In another approach, a cube of stab inoculated water agar was transferred similarly between a sterile slide and sterile coverslip (Villamizar et al. 2021, Rivalier et al. 1932). These were incubated for at least two weeks before they were observed directly, then coverslips were removed for staining and further observation. Furthermore, the fungus was grown in liquid culture in potato dextrose broth shaken at 150 rpm for seven days in an attempt to trigger blastospore production.

### Pathogenicity assays

Branches with EHS crawlers with no outward evidence of fungal growth were collected from hemlock trees in Morgantown, West Virginia. To date, CL has not been confirmed anywhere in the state. Infested branchlets were randomized and distributed evenly into three replicates of two treatments: dipped completely into at least 20 mL of 10^7^ CL conidia in water or dipped into sterile water as a control. To minimize strain-level variability, a composite inoculum consisting of 3 CL strains (ARSEF 14590, VF14, and BAF11) from the three CL-confirmed locations in North Carolina was used for all pathogenicity tests. Treated branchlets (2.5-3.5 cm in length) were fully submerged in inoculum or water for 10 seconds. Branchlets were held at 22°C in petri dishes in separate plastic containers for each treatment. Damp paper towels were provided in containers to maintain high relative humidity for insects and plants. These treatments were monitored daily for 13 days, except on day 9, to assess emergence and mortality. The results of this bioassay were subjected to Kaplan-Meier log-rank survival analysis to test for significant differences in R (v. 4.2.2) using the survival package (Therneau, 2022).

To demonstrate that CL isolated from EHS could reinfect healthy EHS, and thus fulfill Koch’s postulates, the following methods were used. EHS-infested branches were stored at 22°C in a sealed container with a damp paper towel to encourage crawler development. Once sufficient crawlers had emerged after 2-4 days, crawlers were collected for infection assays. Fresh branchlets without EHS were prepared by dipping entirely into a 20 mL solution of 10^7 CL spores(composite inoculum as described above) suspended in water, and set into four petri dishes. Then, 15 emerged crawlers for each of the four dishes were gently moved to each dish using a small, sterile paintbrush. Cadavers from the bioassay were removed 14 days later, surface sterilized with 10% sodium hypochlorite for 10 seconds, and plated on PDA+ST. The resulting isolates were identified through spore and colony morphology and sequencing of the ITS region.

### DNA extraction, PCR and sequencing

DNA was extracted from 7 representative strains isolated from infected EHS crawlers across three sampling locations in Ashe County, North Carolina using the protocol previously described by Macias et al. (2020). This included CL strains ARSEF 14590, BAF11, BAF14, UM11, VF11, VF41, and VF42. For genome sequencing of ARSEF 14590, DNA was extracted using a DNeasy PowerSoil Pro Kit using manufacturer’s protocols. Extracted DNA was stored at −20°C.

To confirm species identities for our strains and build a phylogeny for *Conoideocrella* and close allies, we first targeted sequencing of the universal fungal barcoding gene (the ribosomal internal transcribed spacer region (ITS), which includes ITS1, 5.8S, and ITS2), and the D1–D2 domains of the 28S rRNA gene (28S), for each of 7 CL strains. Primer names, sequences, and PCR programs are listed in SUPPLEMENTARY TABLE 7. PCR reagents and post-PCR clean-up are as described in Macias et al. 2020. The amplified purified products were Sanger sequenced by Eurofins Genomics (Huntsville, AL, USA) using the same primer pairs.

### Genome sequencing and mining

*Conoideocrella luteorostrata* ARSEF 14590 (CLUM14) was sequenced on the Illumina NextSeq 1000 platform at Marshall University Genomics Core Facility (Huntington, WV). Illumina raw reads were assembled using SPAdes v3.15.2 and AAFTF v0.4.1 (Stajich & Palmer, 2019).

Unassembled reads are accessible through the NCBI Sequence Read Archive (SRA) via accession number SRR24939140, BioProject accession number PRJNA980380, and BioSample accession number SAMN35627742. This Whole Genome Shotgun project has been deposited at DDBJ/ENA/GenBank under the accession JASWJB000000000. The version described in this paper is version JASWJB010000000. Nucleotide sequences for RPB1, RPB2, EF1-ɑ, TUB2 and 18S were extracted from the ARSEF 14590 assembled genome using CL strains NHJ 12516 (EF468905, EF468946, and EF468994), CL NHJ 11343 (EF468801), and BCC 9617 (AY624237) reference sequences as BLAST alignment probes. These extracted sequences are deposited separately in NCBI Nucleotide with accessions listed in TABLE 2. Sequences for TUB2 (OR500271) and 18S (OR492260) were not used in tree building but were deposited as well.

**TABLE 1.**
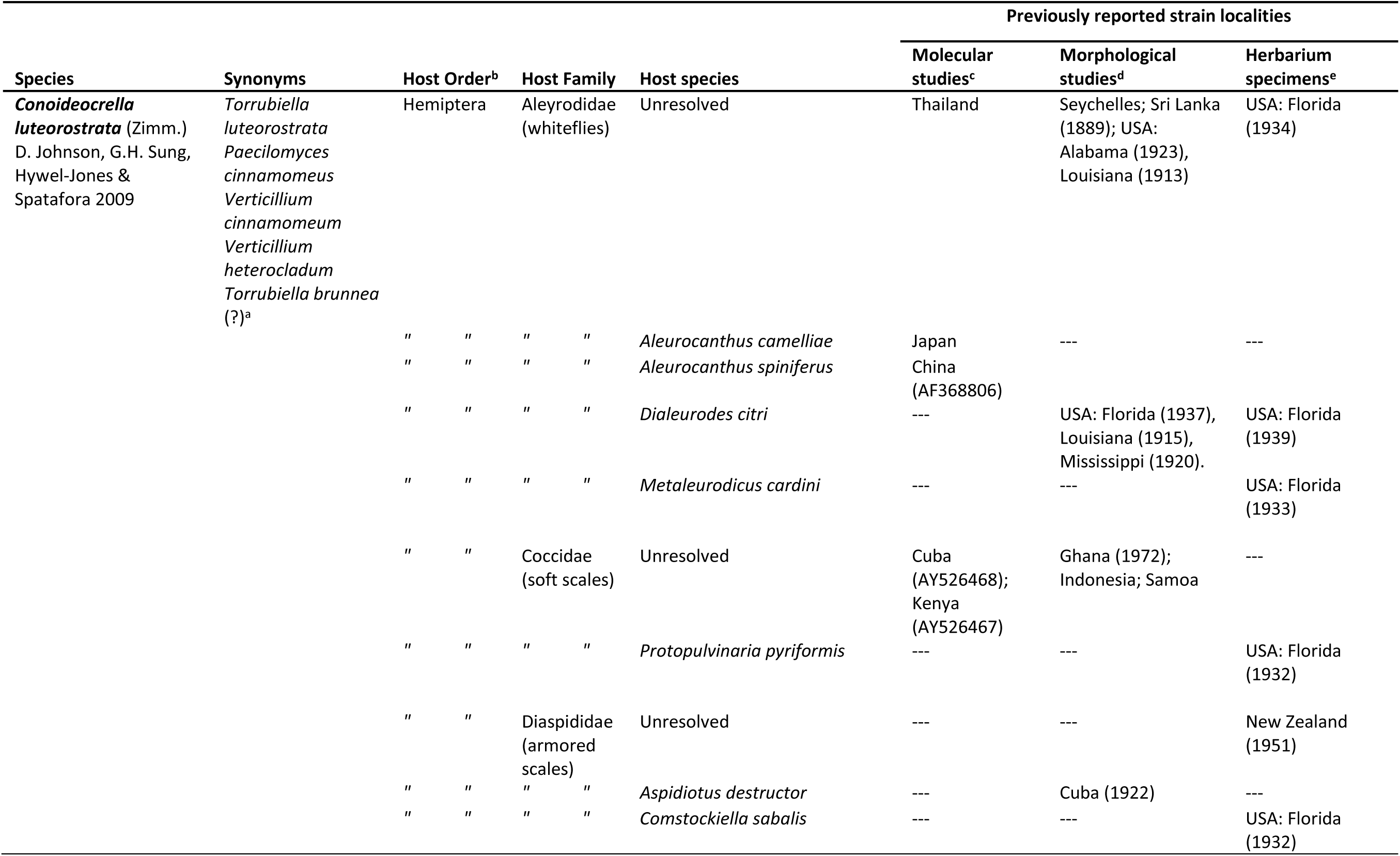
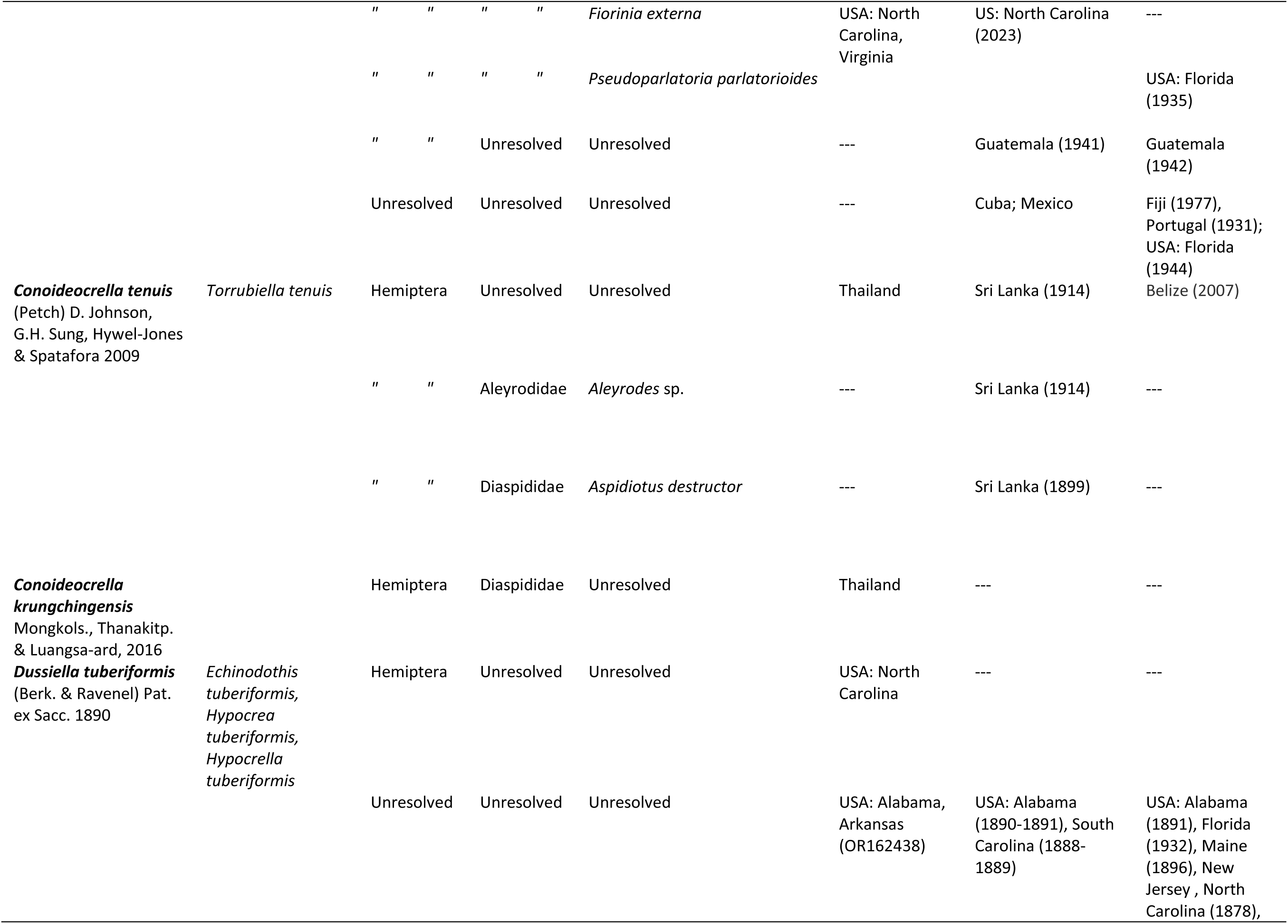
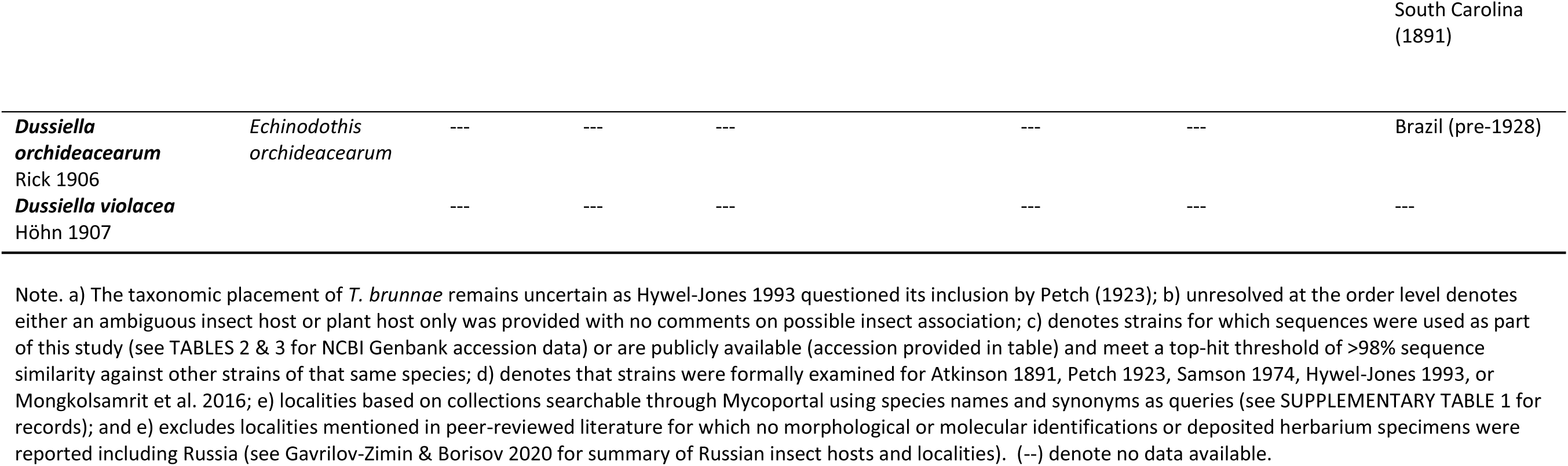
Currently accepted *Conoideocrella* spp. and *Dussiella* spp., host information and locality data based on molecular and morphological data and publicly available herbarium specimens.

**TABLE 2.** Species and isolates used in 54-taxa Clavicipitaceae phylogenetic analysis and their associated metadata.

| Current name<sup>a</sup> | Authority | Synonyms | Strain | 28S | EF1 | RPB1 | RPB2 |
| --- | --- | --- | --- | --- | --- | --- | --- |
| *Aciculosporium*<br>*siamense* | Mongkols., Noisrip. & Luangsa-ard, 2021 | -- | BCC 85384 | MT7430<br>03 | MT7621<br>48 | MT7621<br>49 | -- |
| *Aschersonia*<br>*incrassata* | Mains, 1959 | -- | PC 595 | AY98690<br>9 | AY98693<br>4 | DQ0003<br>35 | -- |
| *Atkinsonella texensis* | (Diehl) Leuchtman & Clay, 1989 | -- | B6156 | KQ06278<br>2 | KQ06273<br>0 | KQ06263<br>0 | KQ06266<br>6 |
| *Balansia henningsiana* | (Möller) Diehl 1950 | -- | GAM 16112 | AY48971<br>5 | AY48961<br>0 | AY48964<br>3 | DQ5224<br>13 |
| *Claviceps fusiformis* | Loveless, 1967 | -- | ATCC 26019 | U17402 | DQ5223<br>20 | DQ5223<br>66 | -- |
| *Claviceps paspali* | F. Stevens & J.G. Hall, 1910 | -- | ATCC 13892 | U47826 | DQ5223<br>21 | DQ5223<br>67 | DQ5224<br>16 |
| *Claviceps purpurea* | (Fr.) Tul. 1853 | *Sphacelia segetum* | GAM 12885 | AF54378<br>9 | AF54377<br>8 | AY48964<br>8 | DQ5224<br>17 |
| *Commelinaceomyces*<br>*aneilematis* | (S. Ito) E. Tanaka, 2020 | -- | MAFF<br>246963 | LC47461<br>7 | LC47462<br>3 | LC47462<br>6 | LC47462<br>9 |
| *Conoideocrella*<br>*luteorostrata* | (Zimm.) D. Johnson, G.H. Sung, Hywel-Jones & Spatafora 2008 | *Torrubiella luteorostrata*, *Paecilomyces cinnamomeus* | ARSEF<br>14590 /<br>UM14 | ON0827<br>55 | OR50027<br>4 | OR50027<br>2 | OR50027<br>3 |
| *Conoideocrella*<br>*luteorostrata* | (Zimm.) D. Johnson, G.H. Sung, Hywel-Jones & Spatafora 2008 | *Torrubiella luteorostrata*, *Paecilomyces cinnamomeus* | NHJ 12516 | EF46884<br>9 | EF46880<br>0 | EF46890<br>5 | EF46894<br>6 |
| *Conoideocrella*<br>*luteorostrata* | (Zimm.) D. Johnson, G.H. Sung, Hywel-Jones & Spatafora 2008 | *Torrubiella luteorostrata*, *Paecilomyces cinnamomeus* | NHJ 11343 | EF46885<br>0 | EF46880<br>1 | EF46890<br>6 | -- |
| *Conoideocrella tenuis* | (Petch) D. Johnson, G.H. Sung, Hywel-Jones & Spatafora 2008 | *Torrubiella tenuis* | NHJ 6293 | EU36904<br>4 | EU36902<br>9 | EU36906<br>8 | EU36908<br>7 |
| *Conoideocrella tenuis* | (Petch) D. Johnson, G.H. Sung, Hywel-Jones & Spatafora 2008 | *Torrubiella tenuis* | NHJ 6791 | EU36904<br>6 | EU36902<br>8 | EU36906<br>9 | EU36908<br>9 |
| *Conoideocrella tenuis* | (Petch) D. Johnson, G.H. Sung, Hywel-Jones & Spatafora 2008 | *Torrubiella tenuis* | NHJ 345.01 | EU36904<br>5 | EU36903<br>0 | -- | EU36908<br>8 |
| *Conoideocrella*<br>*krungchingensis* | Mongkols., Thanakitp. & Luangsa-ard, 2016 |  | BCC 53667 | KJ43507<br>1 | KJ43510<br>0 | -- | -- |
| *Conoideocrella*<br>*krungchingensis* | Mongkols., Thanakitp. & Luangsa-ard, 2016 |  | BCC 36100 | KJ43508<br>0 | KJ43509<br>7 | -- | -- |
| *Dussiella tuberiformis* | (Berk. & Ravenel) Pat. ex Sacc. 1890 | *Echinodothis tuberiformis* | JFW_NC200<br>0 | JQ25700<br>9 | JQ25702<br>7 | JQ25701<br>5 | JQ25702<br>0 |
| *Epichloe typhina* | (Pers.) Brockm. 1863 | -- | ATCC 56429 | U17396 | AF54377<br>7 | AY48965<br>3 | DQ5224<br>40 |
| *Helicocollum krabiensis* | Luangsa-ard, Mongkols., Noisripoom & Thanakitp., 2017 | -- | BCC71372 | KT22232<br>5 | KT22234<br>0 | KT22233<br>3 | -- |
| *Hypocrella nectrioides* | Thaxt., 1921 | -- | GJS 89-104 | U47832 | DQ5223<br>47 | DQ5223<br>93 | DQ5224<br>48 |
| *Hypocrella raciborskii* | Zimm., 1901 | *Aschersonia placenta* | BCC 7869 | EF46907<br>4 | EF46905<br>6 | EF46908<br>5 | EF46910<br>4 |
| *Hypocrella schizostachyi* | Hennings, 1908 | -- | BCC 14123 | DQ5187<br>71 | DQ5223<br>46 | DQ5223<br>92 | DQ5224<br>47 |
| *Hypocrella siamensis* | Hywel-Jones & Mongkols., 2009 Hywel-Jones & Mongkolsamrit 2009 | *Aschersonia badia* | BCC 8105 | DQ5187<br>52 | DQ5223<br>17 | DQ5223<br>63 | DQ5224<br>11 |
| [*Hypocrella*] *hypoxylon* var. *piluliformis* | Berk. & M.A. Curtis, 1892 | *Balansia pilulaeformis*, *Balansiopsis piluliformis*, *Dothidea piluliformis* | AEG 942 | AF54378<br>8 | DQ5223<br>19 | DQ5223<br>65 | DQ5224<br>14 |
| *Keithomyces neogunnii* | (T.C. Wen & K.D. Hyde) Luangsa-ard, Thanakitpipattana & Samson, 2020 | -- | BUM_415 | MH1438<br>28 | MH1438<br>61 | MH1438<br>76 | MH1438<br>91 |
| *Marquandomyces marquandii* | (Massee) Samson, Houbraken & Luangsa-ard, 2020 | *Metarhizium marquandii*, *Paecilomyces marquandii* | CBS 127132 | MT0788<br>57 | MT0788<br>49 | MT0788<br>65 | MT0789<br>22 |
| *Metapochonia gonioides* | (Drechsler) Kepler, S.A. Rehner & Humber, 2014 | *Pochonia gonioides*, *Verticillium gonioides* | CBS 891.72 | AF33955<br>0 | DQ5223<br>54 | DQ5224<br>01 | DQ5224<br>58 |
| *Metarhizium album* | Petch, 1931 | -- | ARSEF 2082 | DQ5187<br>75 | DQ5223<br>52 | DQ5223<br>98 | DQ5224<br>52 |
| *Metarhizium anisopliae* | (Metschn.) Sorokin, 1883 | *Isaria anisopliae*, *Penicillium anisopliae* | ARSEF 3145 | AF33953<br>0 | AF54377<br>4 | DQ5223<br>99 | DQ5224<br>53 |
| *Metarhizium flavoviride* | W. Gams & Rozsypal, 1973 | -- | ARSEF 2037 | AF33953<br>1 | DQ5223<br>53 | DQ5224<br>00 | DQ5224<br>54 |
| *Metarhizium guizhouense* | Q.T. Chen & H.L. Guo, 1986 | *Cordyceps taii*, *Metacordyceps taii*, *Metarhizium taii* | ARSEF 5714 | AF54378<br>7 | AF54377<br>5 | DQ5223<br>83 | DQ5224<br>34 |
| *Moelleriella epiphylla* | (Massee) P. Chaverri & K.T. Hodge 2008 | *Hypocrella epiphylla* | PC386 | EU39258<br>2 | EU39265<br>9 | EU39271<br>0 | -- |
| *Morakotia fusca* | Mongkols., Noisrip., Khons., Thanakitp. & Luangsa-ard, 2021 | -- | BCC 79272 | KY79486<br>1 | KY79485<br>6 | KY79486<br>5 | -- |
| *Myriogenospora atramentosa* | (Berk. & M.A. Curtis) Diehl 1950 | -- | AEG9632 | AY48973<br>3 | AY48962<br>8 | AY48966<br>5 | DQ5224<br>55 |
| *Nigelia martialis* | Luangsa-ard & Thanakitp., 2017 | *Metacordyceps martialis*, *Metarhizium martialis* | BUM_713 | MH1438<br>24 | MH1438<br>58 | MH1438<br>73 | MH1438<br>88 |
| *Orbiocrella petchii* | (Hywel-Jones) D. Johnson, G.H. Sung, Hywel-Jones & Spatafora 2008 | *Torrubiella petchii* | NHJ 5318 | EU36904<br>0 | EU36902<br>1 | EU36906<br>2 | EU36908<br>0 |
| *Orbiocrella petchii* | (Hywel-Jones) D. Johnson, G.H. Sung, Hywel-Jones & Spatafora 2008 | *Torrubiella petchii* | NHJ 6209 | EU36903<br>9 | EU36902<br>3 | EU36906<br>1 | EU36908<br>1 |
| *Orbiocrella petchii* | (Hywel-Jones) D. Johnson, G.H. Sung, Hywel-Jones & Spatafora 2008 | *Torrubiella petchii* | NHJ 6240 | EU36903<br>8 | EU36902<br>2 | EU36906<br>0 | EU36908<br>2 |
| *Papiliomyces* sp. YZ-2021a | N/A | -- | YC20061404 | MZ702102 | MZ955881 | MZ955877 | OM419144 |
| *Parametarthizium hingganense* | S. Gao, W. Meng, Li Xiang Zhang, Q. Yue, L. J. Xu, 2021 | -- | SGSF355 | MN061635 | MN065770 | MN917170 | MT939494 |
| *Periglandula ipomoeae* | U. Steiner, Leistner & Schardl, 2011 | -- | IasaF13 | AFRD01000776 | AFRD01000133 | AFRD01000003 | AFRD01000102 |
| *Pochonia chlamydosporia* | (Goddard) Zare & W. Gams, 2001 | *Cordyceps chlamydosporia, Diheterospora chlamydosporia, Verticillium chlamydosporium* var. *chlamydosporium*, *Verticillium catenulatum*, *Metacordyceps chlamydosporia* | CBS 101244 | DQ518758 | DQ522327 | DQ522372 | DQ522424 |
| *Pochonia cordycepsociata* | H. Huang, M. Wang & L. Cai, 2015 | -- | CGMCC 317365 | KM263573 | KM263584 | KM263576 | KM263579 |
| *Pseudogibbellula formicarum* | (Mains) Samson & H.C. Evans, 1973 | -- | BCC 84257 | MT512653 | MT533480 | MT533473 | -- |
| *Purpureomyces pyriformis* | Luangsa-ard, Noisripoom, Himaman, Mongkolsamrit, Thanakitpipattana & Samson, 2020 | -- | BCC 85348 | MN781871 | MN781728 | MN781773 | MN781820 |
| *Regiocrella camerunensis* | P. Chaverri & H.C. Evans, 2006 | -- | ARSEF 7682 | DQ118735 | DQ118743 | DQ127234 | -- |
| *Rotiferophthora angustispora* | (G.L. Barron) G.L. Barron 1991 | -- | CBS 101437 | AF339535 | AF543776 | DQ522402 | DQ522460 |
| *Samuelsia chahalensis* | P. Chaverri & K.T. Hodge, 2008 | -- | PC560 | EU39263 | EU392691 | EU392743 | -- |
| *Shimizuomyces paradoxus* | Kobayasi, 1981 | -- | EFCC 6279 | EF469084 | EF469071 | EF469100 | EF469117 |
| *Tolypocladium ophioglossoides* | (Ehrh. ex J.F. Gmel.) Quandt, Kepler & Spatafora, 2014 | *Cordyceps ophioglossoides, Elaphocordyceps ophioglossoides* | OSC 106405 | AY489723 | AY489618 | AY489652 | DQ522429 |
| *[Torribiella] hemipterigena* | Petch, 1931 | *Verticillium hemipterigenum* | Allpaths-LG | CDHN01000016 | CDHN01000003 | CDHN01000002 | CDHN01000003 |
| *Ustilaginoidea virens* | (Cooke) Takah., 1896 | *Claviceps virens, Villosiclava virens* | ATCC 16180 | -- | JQ257026 | JQ257014 | JQ257019 |
| *[Verticillium] epiphytum* | Hansford, 1943 | *Verticillium epiphytum* | CBS 384.81 | AF339547 | DQ522361 | DQ522409 | DQ522469 |
| *Yosiokobayasia kusanaginensis* | (Kobayasi & Shimizu) Samson, Luangsa-ard & Thanakipattana, 2020 | *Metacordyceps kusanaginensis, Metarthizium kusanaginense* | TNS-F 18494 | JF415972 | JF416014 | JN049890 | -- |

Note. a) Square brackets ([ ]) around a genus indicates that the name awaits appropriate action by the research community to be transferred to another genus as indicated in NCBI Taxonomy Browser. (--) denote no data available.

**TABLE 3.**
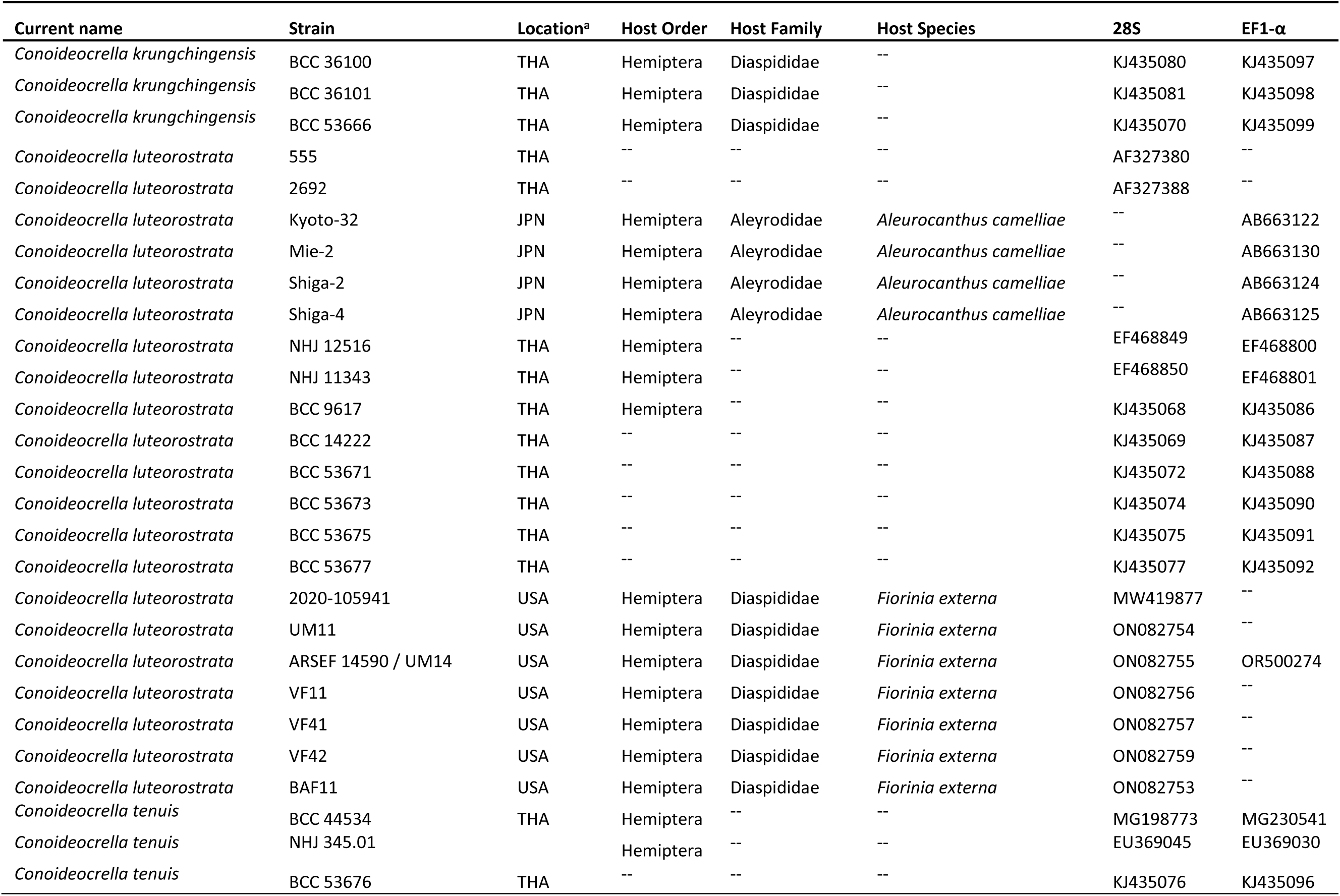
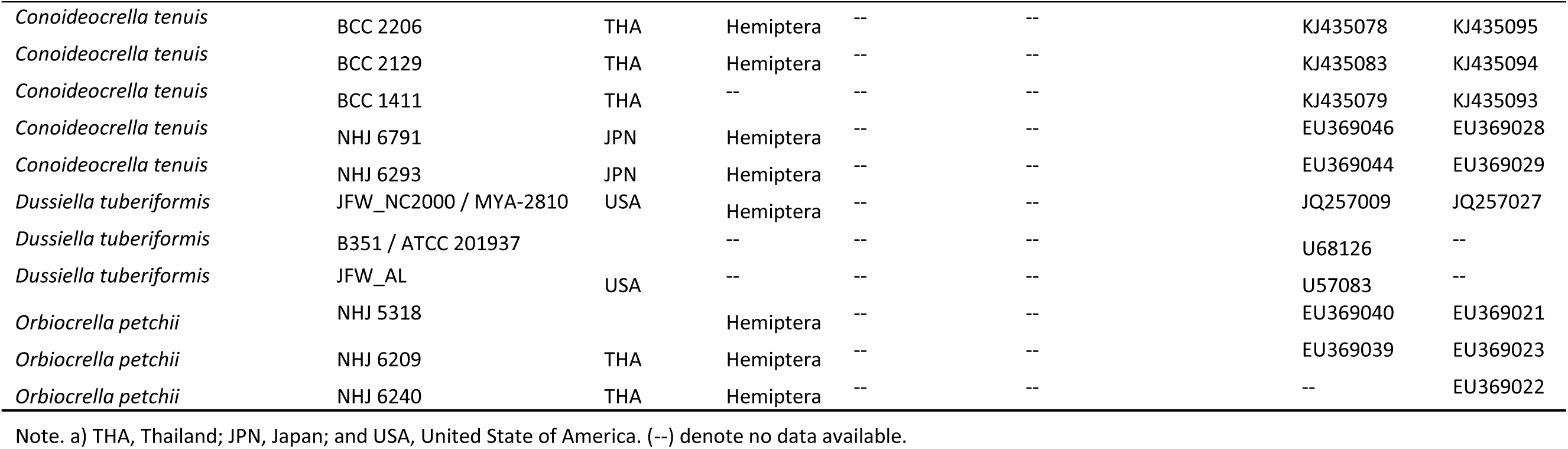
Species and isolates used in 38-taxa *Conoideocrella*-focused phylogenetic analysis and their associated metadata.

### Alignments, model selection and phylogenetic analyses

Chromatograms for Sanger sequences were quality-checked using default parameters, clipped, and manually corrected in Geneious Prime (2023.1.2). Two datasets were compiled: 1) a 54-taxa 4-locus (RPB1, RPB2, EF1, 28S) dataset (hereafter referred to as the Clavicipitaceae dataset) representing the diversity of the family based on previous phylogenetic studies (Johnson et al. 2009, Kepler et al. 2012, and Mongkolsamrit et al. 2016), and 2) a 38-taxa 2-locus *Conoideocrella*-focused (EF1, 28S) dataset (hereafter referred to as the CF dataset) informed by the topology of the 4-locus analysis and previous phylogenetic studies (Artjariyasripong et al., 2001; Luangsa-ard et al., 2004; Saito et al., 2012; Urbina & Ahmed, 2022). Both datasets included novel sequence data for CL ARSEF 14590. The CF dataset also included 28S sequence data for five additional CL strains. Strain tables for each analysis are available in TABLE 2 (Clavicipitaceae dataset) and TABLE 3 (*Conoideocrella*-focused dataset).

Each locus was aligned separately using MAFFT (Katoh & Standley, 2013) on the GUIDANCE2 server (http://guidance.tau.ac.il/; Landan & Graur, 2008; Sela et al., 2015), and individual residues with GUIDANCE scores <0.5 were masked. An intron in RPB1 (positions 118 to 247bp) was deleted. The best nucleotide substitution models for each locus were selected by ModelTest in MEGA11 (Tamura et al. 2021), using the corrected Akaike information criterion (AlCc) score. GTR+G was determined to be the optimal model for all analyses.

Single-locus alignments and concatenated alignments for both datasets were all used for tree building. Maximum Likelihood (ML) trees were created using RAxML v.8.2.12 (Stamatakis, 2014) with 1000 bootstrap replicates. Bayesian Inference (BI) trees were generated using MrBayes 3.2.7a (Ronquist et al., 2012), using 1M generations except for the 28S Clavicipitaceae analysis which required 3.4M generations for the standard deviation of split frequencies to fall below 0.01. For BI analyses, one cold chain and three heated chains were used for each run, and the first 25% of generations were discarded as burn-in. Finally, runs were checked for convergence in Tracer 1.7.1 (Rambaut et al., 2018). For concatenated datasets, the ML tree was used for topology and branch lengths, and was annotated with BI support values where topology was in agreement. Trees were viewed and prepared for publication using FigTree 1.4.4 (Rambaut et al., 2018) and Inkscape 0.92.2 (https://www.inkscape.org/). Resulting trees, alignments, and other data are available on GitHub: https://github.com/HanaBarrett/EHS-CL-Analysis.

## Results

### Morphological investigation

An emergent fungal epizootic of *Conoideocrella luteorostrata* (CL) was found impacting the first instar crawler stage of elongate hemlock scale (EHS) on planted Fraser fir trees across three Christmas tree farms in Ashe County, North Carolina in the summer of 2020. Hundreds of thousands of mycosed individuals were observed across the three NC sampling locations with the heaviest infections occurring at VF, followed by UM then BAF (FIG. 1A-B). No mycosed EHS were found in Virginia at MRO, despite the presence of EHS at this location. Neither healthy nor mycosed EHS were recovered from MR, though healthy Fraser fir needles were sampled. At VF, CL was also observed and isolated from one adult female EHS. Infected EHS were identifiable by the distinct orange mat of hyphae (stromata) covering the nymph cadavers. Mycosed EHS were abundant, with multiple infected cadavers on each affected needle (FIG. 1A).

Morphological studies were undertaken to compare four CL strains isolated from two sampling sites (UM, VF) in North Carolina with previously reported measurements for this species (Samson, 1974; Hywel-Jones, 1993; Saito et al., 2012), other *Conoideocrella* species (Hywel-Jones, 1993, Mongkolsamrit et al., 2016) and *Dussiella tuberiformis* (Atkinson, 1891).

Newly established CL colonies were a distinct tannish orange color that gradually changed to cinnamon-brown on PDA+ST. After about a week of growth, CL colonies begin to secrete a yellow soluble pigment (FIG. 1G-H)t, eventually turning a purplish red as the cultures aged (FIG. 1F). This was apparent in both the agar and as guttation droplets atop the older inner mycelium (FIG. 1G; right panel). These droplets were also observed atop CL colonies on Czapek-Dox agar (photos not shown). After several weeks, colonies were covered with green conidia (FIG. 1E-F).

Subsequent microscopic examination of these four CL strains in lactic acid (+/-cotton blue) confirmed the presence of whorled (verticillate) conidiophores with flask-shaped phialides and hyaline, smooth-walled, aseptate conidia with acute ends (FIG. 1I-L). As cultures aged, basipetal chains of conidia were observed resulting in a characteristic powdery, green appearance (FIG. 1F,J). Based on the taxonomic keys of Samson (1974) and subsequent descriptions from Saito et al. (2012), the fungus aligned morphologically with *Paecilomyces cinnamomeus*, the asexual state of CL.

Conidia averaged 6.9 (5.4-8.0) µm in length x 2.6 (2.0-3.3) µm in width across the four examined CL strains (TABLE 4). Among them, mean spore lengths and widths were significantly larger (p<0.05) for two strains. ARSEF 14590 (mean 7.4 x 2.5 µm) and VF11 (mean 7.2 x 2.8 µm) had significantly larger mean conidial lengths (p<0.05) compared to UM11 (mean 6.6 x 2.7 µm) and VF41 (mean 6.5 x 2.4 µm) (FIG. 2). For conidial width, VF11 (7.2 x 2.8 µm) has significantly larger spore widths (p<0.05) compared to VF41 (6.5 x 2.4 µm), but neither were significantly different from UM11 or ARSEF 14590. Overall, our results suggest that there is considerable variation in conidial length among our isolates. However, the conidia lengths for all four strains fell within the ranges reported by Hywel-Jones (1993), Samson (1974) and Saito et al. (2012). Additionally, all four CL strains had conidial widths that overlapped with Hywel-Jones (1993). UM11 and VF41 also had conidial widths that overlapped with Saito et al. (2012) and VF41 overlapped with Samson (1974) (FIG. 2). Raw spore measurements are available in SUPPLEMENTARY TABLE 5. Nearly all strains examined including measurements reported from aforementioned taxonomic papers overlapped partially with the conidia measurements reported for a hirsutella-like asexual state of CK (Mongkolsamrit et al., 2016), which covered a much larger range of both length and width. However, unlike CK, CL conidia were exclusively aseptate. Measurements for *Dussiella tuberiformis* were distinct from all CL strains, with the exception of minimal overlap in spore length in ranges reported by Hywel-Jones (1993) (FIG. 2). Ascospore measurements were not compared, as the sexual stage was not observed on mycosed EHS during this study: perithecia have never been reported from U.S. hemipterans infected with CL, though herbarium specimens of CL from AL, FL, LA, and MS have not been examined for such fruiting bodies.

**TABLE 4.** Ascospore and conidia measurements for *Conoideocrella* species and *Dussiella tuberiformis* with associated references.

| Species ID | Ascospores (um) | Asexual state | Conidia (um) | N (isolates / spores)<sup>a</sup> | Reference |
| --- | --- | --- | --- | --- | --- |
| *Conoideocrella luteorostrata* | 460-590 x 1.5-2 | *Paecilomyces cinnamomeus* | 6.5-8.3 x 2.0-3.6 | -- | Hywel-Jones 1993 |
|  | -- | " " | 5.2-7.5 x 1.7-2.5 | -- | Samson 1974 |
|  | -- | " " | 5.2-7.6 x 2.3-2.6 | 12 / 600 | Saito et al. 2012 |
|  | -- | " " | 6.2 (4.5-8.0) x 1.9 (1.8-2.9) | 4 / 100 | This study |
| *Conoideocrella tenuis* | 380-500 x 1.5-2 | unknown | -- |  | Hywel-Jones 1993 |
| *Conoideocrella krungchingensis* | (80-)110-150 x 1.5-2.5 | hirsutella-like | 8-15 x 2-4 | -- | Mongkolsamrit et al. 2016 |
| *Dussiella tuberiformis* | 450-750 x 14 | sphacelia-like | 7-10 x 3.5-4 | -- | Atkinson 1891 |

Note. A) Denotes total isolates used and total spores measured across all isolates. (--) denote no data available.

A hirsutella-like asexual state with paired aseptate to uniseptate conidia formed on simple conidiophores was also observed for ARSEF 14590 after 14 days (FIG 3). A mucous sheath was visible immediately surrounding the paired fusiform to slightly curved spores atop each conidiophore, as evidenced by the absence of mountant infiltration (FIG 3B-E). Sporulation was sparse, spores stuck together and were and intermixed with conidiophores of the paecilomyces-like synanamorph, which limited the ability to report measurements specifically for this synanamorph except for uniseptate spores, which were completely absent from the paecilomyces-like colonies examined on standard growth media and conditions (FIG 1I-L). A single uniseptate spore in mucous sheath and still attached to its conidiophore measured 8.9 x 2.0 µm, exceeding the length of all previously reported and measured conidia across all CL strains. No propagules or blastospores were observed in liquid culture.

**Figure 3.**
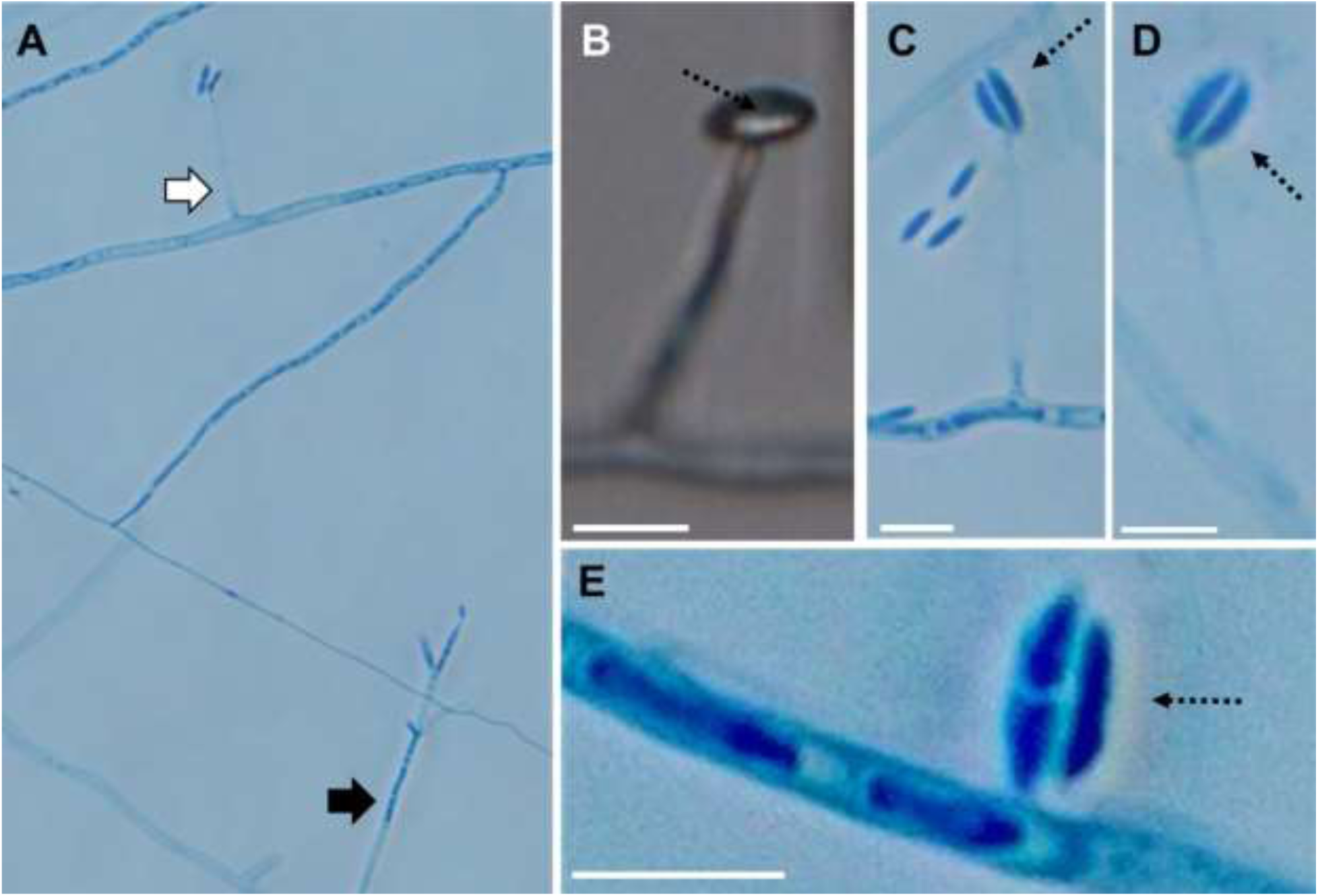
Conidiophores and conidia of hirsutella-like asexual stage of *C. luteorostrata* strain ARSEF 14590 / CLUM14 sporulating beneath glass slipcovers on stab-inoculated PDA slides. A) Both paecilomyces-like (black arrow) and hirsutella-like (white arrow) asexual states co-occurring on the same slide, B-D) conidiophores showing paired terminal fusiform conidia with mucous sheath (dashed arrows show areas absent of mountant infiltration), E) a transversely septate and aseptate conidium still paired within a mucous sheath detached from the conidiophore. Scale bars in B-E, 10 µm.

**Figure 4.**
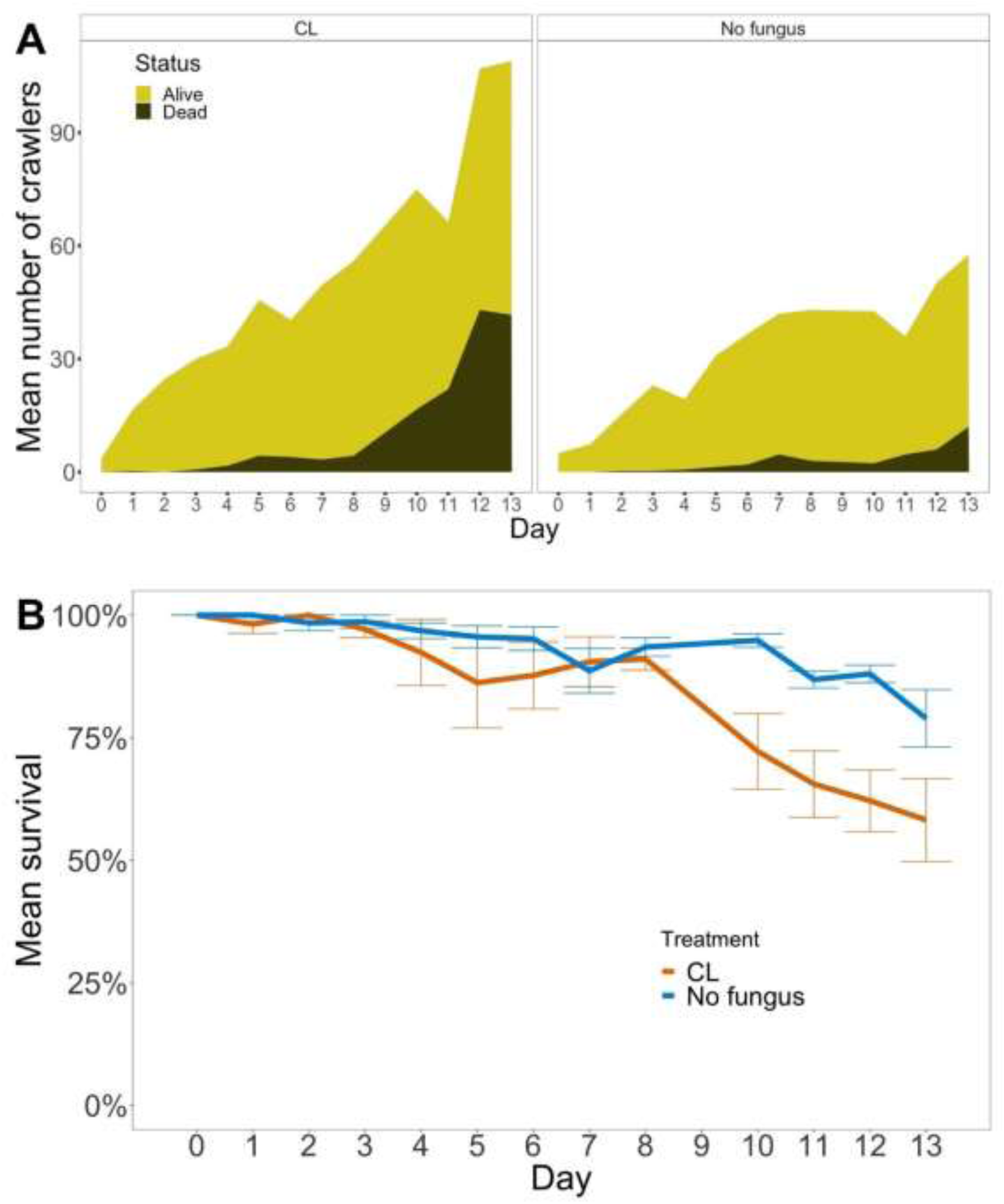
Crawler emergence by treatment over the course of our bioassay (A). Mean total crawlers (across 3 replicates) are represented by the yellow area whereas mean dead crawlers are represented in black. Calculated percent survival for each treatment (B).

For temperature growth assays, our two examined CL strains grew optimally around 22°C, with an estimated radial growth rate of 0.95 mm/day on PDA+ST, but neither actively grew at 10°C or 30°C. Because some plates developed satellite colonies over the course of the first two week period, only 4 plates were available per isolate per temperature, with the exception of BAF45 that had 3 at 10°C and 5 at 22°C. Both strains were able to resume normal growth when returned to 22°C after 7 days exposure to either temperature.

### Culture-based survey of fir needles for CL

A total of 82 fungal isolates representing 31 fungal morphotypes were recovered from 60 Fraser fir needles (38 living and 12 dead) sampled across five sampling locations. 24 total and 18 unique morphotypes were recovered from living fir needles. 14 total morphotypes were recovered from EHS + CL+ trees, of which 6 morphotypes were absent from EHS+ CL- and EHS-CL-trees. However, none of the 31 recovered morphotypes, including the 6 unique morphotypes associated with CL+ trees, shared any characteristics that distinguish CL */* PC.

### Superficial arthropod survey for CL-infected arthropods

Seventy non-sessile arthropods were collected from our CL+ branch samples (SUPPLEMENTARY TABLE 4). No outward fungal infections were observed in these arthropods. The most common arthropod encountered was the beetle *Cybocephalus nipponicus*, a known predator of EHS, which was collected from VF and BAF: both of these sites were not being managed for production during collection, and, as such, both hosted denser EHS populations. Two parasitoid wasps were also collected: one of these was observed ovipositing into an EHS adult female prior to collection. We identified these parasitoid wasps as *Encarsia citrina*, a known parasitoid of EHS. EHS-parasitoid wasps were also observed developing within the tests of collected female elongate hemlock scales under microscopic examination. None of these natural enemies of EHS or other arthropods exhibited outward fungal infections. No isolations were attempted from these outwardly asymptomatic insects.

### Laboratory pathogenicity bioassays

In our first bioassay, EHS-infested needles were treated with a CL suspension, and crawlers were allowed to emerge over the course of the bioassay. EHS are known to have continuous emergence across the season, and this was observed for the duration of both bioassays. This presented challenges for counting moving crawlers when numbers rose above ∼20 individuals, and it also resulted in continuous exposure to CL: some crawlers were initially infected on day 0, while others were infected on day 13 (FIG. 4A). This continuous emergence resulted in percent survival increasing on some days (FIG. 4A). Crawler vitality was recorded daily for both treatments for Kaplan-Meier survival analysis. CL-treated EHS and controls were significantly different (log-rank test, p = 0.03). The survival curve (FIG. 4B) documenting daily percent survival in this bioassay revealed lower survival in the CL treatment late in our bioassay (i.e., days 10-13), but continuous crawler emergence prevented survival from falling below 50%.

In our second bioassay, crawlers were introduced onto CL-treated branchlets and cadavers were collected two weeks after exposure. An asymptomatic crawler, a treated crawler exhibiting melanization (insect immune response to infection) and fungal outgrowth, and a cadaver with visible conidiophores with paired to whorled phialides typical of CL can be seen in FIG. 5A-C. CL was isolated from three of five surface-sterilized crawler cadavers and identified through colony and conidia morphology and Sanger sequencing of the ITS region. CL was not isolated from any of three surface-sterilized crawlers randomly sampled from the control group.

**Figure 5.**
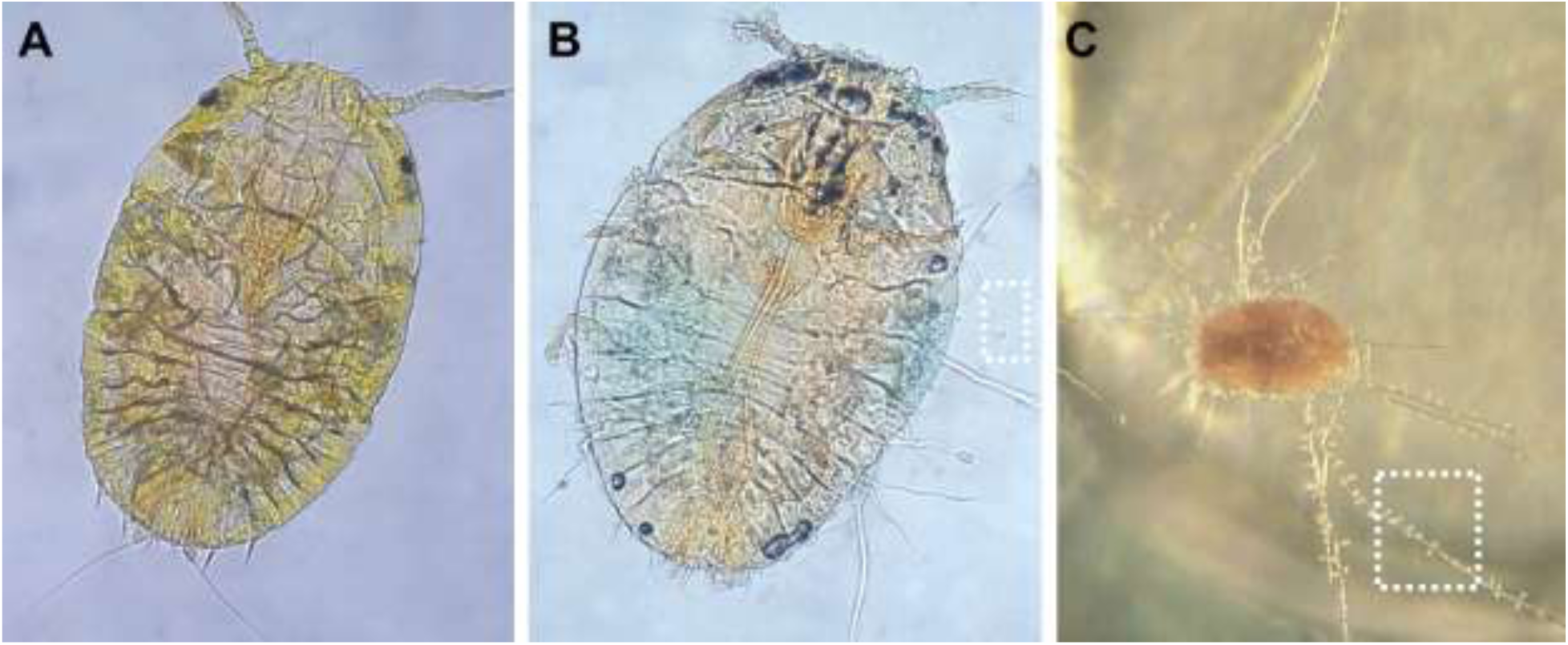
Asymptomatic EHS crawler from control treatment at 40x magnification (A) compared to crawler from CL+ treatment (40x) exhibiting melanization, which was associated with infection (B). Dead mycosed crawler with *C. luteorostrata* outgrowth under dissecting microscope (C). Dashed white boxes in B and C emphasize fungal growth with obvious paired to whorled conidiophores characteristic of CL seen in C.

**Figure 6.**
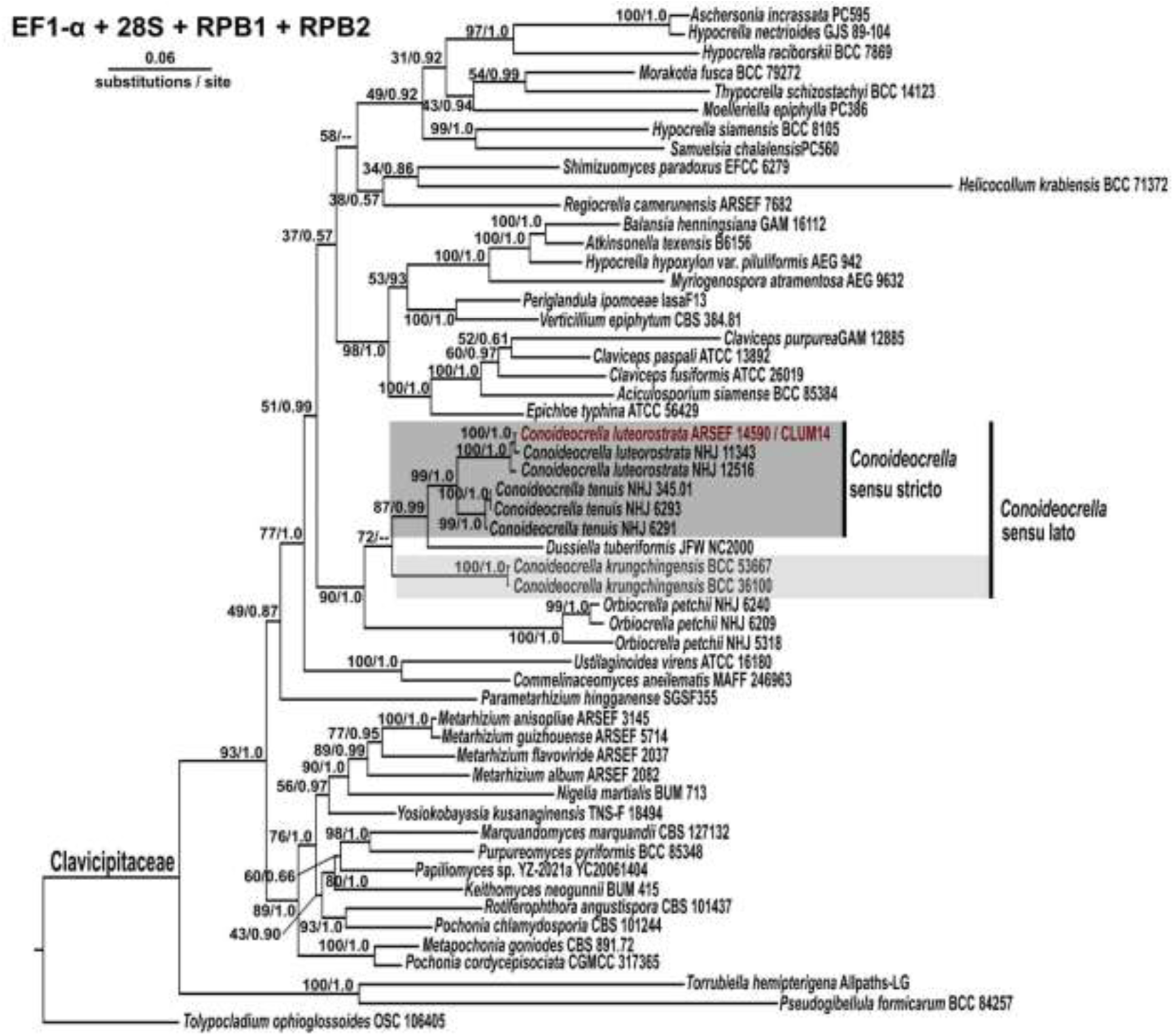
Concatenated phylogenetic tree for the Clavicipitaceae dataset (28S + EF1-α + RPB1 + RPB2) dataset. Topology and branch lengths shown are from the ML analysis. Bootstrap support and posterior probabilities are indicated for each node supported in the ML analysis (ML/BI). Strain metadata including GenBank accession numbers are listed in Table 2.

**Figure 7.**
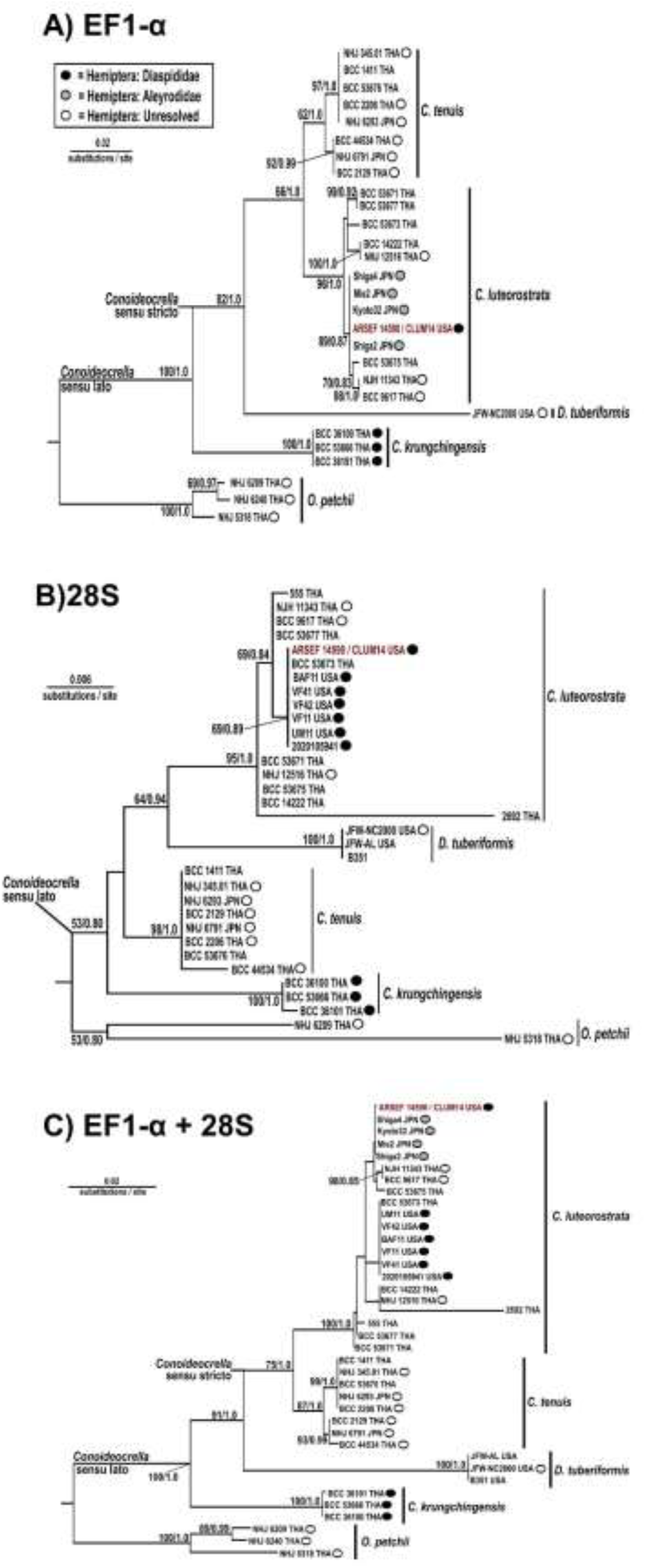
Phylogenetic trees for EF1-α (A), 28S (B), and concatenated (C) *Conoideocrella*-focused datasets. Topology and branch lengths shown are from the ML analysis. Bootstrap support and posterior probabilities are indicated for each node supported at or above 50% in the ML analysis (ML/BI). Country of origin and insect host associations are denoted for all taxa where data are available with key for host taxa in tree 6A. Strain metadata including GenBank accession numbers are listed in Table 3.

### CL molecular identification

BLASTN searches against NCBI GenBank using our isolates as queries revealed three of our seven strains (ARSEF 14590, UM11, and VF11) were 100% identical to two CL ITS sequences (MW419875, MW419876) that were recently deposited from mycosed EHS in North Carolina (independent of our study; Urbina & Ahmed, 2022) and one CL ITS sequence (AB649298) from a mycosed whitefly in Japan in 2009 (Saito et al., 2012). To identify the degree of sequence similarity among the 7 novel CL isolates, pairwise BLASTN searches were conducted and revealed sequence similarity ranging from 99.81 – 100% across all ITS sequences (deposited under GenBank accessions ON081993-ON081999). BLASTN searches were also conducted for ARSEF 14590 beta-tubulin 2 (BTUB) and 18S sequences, revealing 99.4% sequence similarity to TUB2 sequences deposited for CL Kyoto-26 (AB663108) and Shiga-4 (AB663113) from Japan, and 100% sequence similarity to 18S sequences deposited for CL NHJ 11343 (EF468995) and CL NHJ 12516 (EF468994) from Thailand.

### Phylogenetic analysis of *Conoideocrella* spp. and closely allied fungi

To examine phylogenetic relationships among diverse members of the Clavicipitaceae, the three formally recognized *Conoideocrella* spp., and multiple CL strains including our seven strains from North Carolina, phylogenetic analyses were conducted for a 54-taxa, 4-locus Clavicipitaceae dataset and separately for a 38-taxa, 2-locus *Conoideocrella-*focused (CF) dataset representing the diversity of strains in *Conoideocrella* and closely allied genera. Single-locus and combined datasets were analyzed for both datasets using two methods of phylogenetic inference for a total of 10 analyses for the Clavicipitaceae dataset and 6 analyses for the CF dataset.

For the Clavicipitaceae dataset, both ML and BI resolved CL as a strongly-supported (>94% bootstrap support / >0.99 posterior probability) monophyletic lineage for all single-locus (RPB1, RPB2, EF1-α and 28S) analyses and in the concatenated 4-locus analysis (FIG. 6, SUPPLEMENTARY FIG. 1-4). CT also received strong support both in the RPB1 and RPB2 single-locus analyses (SUPPLEMENTARY FIG. 3-4) and combined analysis (FIG. 6). CT received moderate (71% / 0.98) support in the EF1-α analysis (SUPPLEMENTARY FIG. 1). RPB1 and RPB2 sequence data was not available for CK, but the species was strongly supported (100% / 1.0) in the 28S, EF1-α, and combined analyses (SUPPLEMENTARY FIG. 1-2).

*Dussiella tuberiformis* formed a well-supported clade with CL and CT in the RPB1, RPB2, and concatenated analyses for the Clavicipitaceae dataset (FIG. 6, SUPPLEMENTARY FIG. 3-4). However, EF1-α and 28S single-locus trees did not resolve congruent topologies (SUPPLEMENTARY FIG. 1-2). In both trees, the relationships among CT, CL, CK and DT could not be resolved, with little to no support in the backbone of both trees. Regardless, no tree containing both CK and DT supports the monophyly of *Conoideocrella*. In the concatenated analysis, DT emerges between the clade that contains CL and CT, and the CK clade (FIG. 6).

*Orbiocrella petchii* (OP) formed a well-supported, genealogically-exclusive lineage across all analyses, with the exception of the 28S analysis, which revealed OP to be paraphyletic with one strain grouping with CT (SUPPLEMENTARY FIG. 2). OP formed a moderately to well-supported clade with CL, CT and DT in the RPB1, RPB2, and concatenated trees (FIG. 6, SUPPLEMENTARY FIG. 3-4). However, EF1-α and 28S single-locus trees were incongruent: placement of OP occurred elsewhere as a single clade (EF1-α; SUPPLEMENTARY FIG. 1) or as paraphyletic (28S; SUPPLEMENTARY FIG. 2).

For the Clavicipitaceae dataset, RPB1 and RPB2 supported the largest number of nodes at ≥ 70% ML / >0.70 BI (n = 23 and 18, respectively). By contrast, 28S was the least informative with 9 supported nodes (SUPPLEMENTARY FIG. 2). Low support in the backbone of all single locus trees emphasizes the need for multi-locus phylogenetic studies. Given the paucity of publicly available C*onoideocrella* sequence data for RPB1 and RPB2 (n = 5 / locus including ARSEF 14590), only 28S and EF1-α were advanced for the *Conoideocrella*-focused (CF) 38-taxa dataset. The results from the Clavicipitaceae dataset also helped inform appropriate ingroup and outgroup taxa for the CF dataset, with *Conoideocrella* and *Dussiella* serving as the ingroup and *Orbiocrella* as the outgroup.

For the CF dataset, both ML and BI resolved CL as a strongly-supported (>95% / > 0.99) monophyletic lineage for both the single-locus analyses (EF1-α and 28S) and the concatenated analysis (FIG. 7). The EF1-α single locus analysis further resolved one well-supported (89% / 0.87) clade within CL containing our isolate (ARSEF 14590), four Japanese strains from whitefly, and three strains from Thailand (FIG. 7A). The 28S analysis resolved a moderately-supported (69% / 0.84) clade in CL that contained ARSEF 14590, the same three strains from Thailand, and additional strains not present in the EF1-α tree due to missing data, including more North Carolina strains from this study and strain 2020-105941, an independently acquired strain from elongate hemlock scale in Virginia (FIG. 7B). The Japanese strains from whitefly were also excluded from the 28S analysis, as no sequence data was available. Any fine-level resolution among CL isolates supported by single-locus trees was absent in the concatenated tree (FIG 7C).

CT and CK also formed well-supported genealogically exclusive lineages across all analyses except for CT in the EF1-α analysis, which received only moderate (62% / 1.0) support (FIG. 7). Despite the moderate support for CT, EF1-α and concatenated analyses further resolved two clades within CT, both containing strains from Thailand and Japan (FIG. 7A, 7C).

Despite the strong support for all three individual species from both methods of phylogenetic inference, C*onoideocrella* is polyphyletic (FIG. 7)*. Dussiella tuberiformis* formed a strongly supported genealogically exclusive lineage sister to either CL (28S: 64% / 0.94), or sister to a clade containing both CL and CT (82% / 1.0 for EF1-αand 91% / 1.0 for the concatenated tree). In both the single-locus and concatenated trees, CK fell outside the clade containing CL, CT, and *D. tuberiformis* (FIG. 7).

For the CF dataset, EF1-α had a higher number of supported nodes (12 nodes at ≥ 70% / 0.7), compared to 28S (4 nodes at ≥ 70% / 0.7). The combining of EF1-α and 28S datasets weakens intraspecies resolution. Based on the results of the single-locus and 2-locus datasets, EF1-α alone appears to be the most robust candidate locus for intraspecies-level resolution for both CL and CT,providing strong support along the backbone of the phylogeny.

## Discussion

Our recent discovery of a naturally occurring epizootic of the anamorphic state of *C. luteorostrata* on elongate hemlock scale in North Carolina Christmas tree farms suggests potential for this fungus as a biocontrol agent. This exotic pest, though economically significant, does not currently have a commercially available fungal biocontrol option for tree farms, where it primarily infests fir. As with many armored scale insects, chemical insecticides may be ineffective due to varying non-target effects on natural predators, insufficient spray coverage due to tree size or timing of application, and /or diminished efficacy due to natural resistance of adult females (Sidebottom, 2016). A 2022 survey by Urbina & Ahmed supports the relatively recent expansion of this fungus across EHS populations impacting Christmas tree production in VA, NC, and MI, but more expansive surveys across other plant and insect hosts are needed, especially given the numerous historic reports of anamorphic CL from whiteflies, soft scales and armored scales, including citrus whitefly across at least four southeastern U.S. states between 1913-1939 (TABLE 1). CL co-occurs with *Dussiella tuberiformis* and at least two other Clavicipitaceae whitefly entomopathogens, *Aschersonia aleyrodis* and *A*. *goldiana* (Fasulo & Weems, 1999; Martini et al., 2022), across much of the same geographic area. Dubious iNaturalist observations coupled with the limited host data for these species, adds further uncertainty to host and species boundaries for these fungi across the southeastern U.S.

In this study, we expanded on the current understanding of the U.S. population of CL with a robust phylogenetic and morphological investigation of our strains and compared them to CL, other *Conoideocrella* spp. and DT strains from the peer-reviewed literature. Our work confirmed that CL strains recovered from North Carolina (including strains independently isolated by Urbina & Ahmed (2022)) form a well supported clade within CL that included whitefly strains from Japan and strains from Thailand (FIG. 7A). Our morphological data overlaps considerably with all previously reported measurements for CL, with the exception of conidia widths reported by Hywel-Jones (1993), which cover a wider range of measurements. In addition, a hirsutella-like synanamorph recovered from ARSEF 14590 yielded uniseptate spores that exceeded the spore length maximum for all previously measured CL strains. Host associations for a majority of isolates were taxonomically unresolved below family-level, so it remains unclear if there are any biologically significant patterns across CL and *Conoideocrella*. Consistent with previous literature (Saito et al., 2012), we found non-sessile crawlers are more vulnerable to infection, which may present challenges for identifying insects based on diagnostically relevant morphological characteristics often described from adults (Urbina & Ahmed, 2022). Even under ideal conditions for identifying the host (i.e., when adults are infected), CL quickly envelops and significantly degrades hosts, making their morphological identification below family level impractical (Hywel-Jones 1993).

CL monophyly and its relationship with CT was well-supported in both the larger Clavicipitaceae dataset and the *Conoideocrella*-focused dataset, but none of our analyses support the monophyly of *Conoideocrella*, with various and incongruent placement of CK and DT. The exclusion of DT from the recent taxonomic description of CK resulted in a well-supported, but incomplete perspective of the genetic relationships among the three species of *Conoideocrella*. Both the historic and current morphological data generated for this study provide further insights, some of which further support our phylogenetic conclusions: CK has noticeably different ascospore lengths compared to CL, CT, and DT according to the literature (TABLE 4). On the other hand, CK and CL both produced hirsutella-like anamorphs with occasionally-(CL) to exclusively-(CK) septate conidia singly or in pairs on unbranched short conidiophores. Though this synanamorph was only produced by CL under special circumstances, it produced mostly aseptate conidia similar in size to those reported for the paecilomyces-like synanamorph seen in all examined CL strains examined strains. DT also produces a similar hirsutella-like or sphacelia-like asexual state with aseptate to occasionally uniseptate conidia on “needle-shaped” conidiophores (Atkinson 1891). Despite the presence of a hirsutella-like anamorph in our North Carolina CL strains, the paecilomyces-like synanamorph with aseptate conidia on verticillate conidiophores was the dominant anamorph and observed in all examined strains. Whether CL, CT and DT, or all *Conoideocrella* spp. including CK and DT, should be synonymized as *Dussiella* spp. remains unclear without additional sequence data, conidiophore and conidia observations and measurements for CT. These are currently lacking, as are morphological and molecular datasets for a CL-like fungus that was commonly reported on various insect hosts across the southeastern U.S. in the first half of the last century (see TABLE 1). To tackle these issues, future researchers investigating fungal epizootics on Hemiptera should retain sufficient materials for morphological investigations, DNA sequencing including EF1-ɑ and other protein coding genes including RPB1 and RPB2, and depositing voucher specimens in herbaria for future independent examination. The paucity of sequence data for *Dussiella* will require targeted sequencing of strains from culture collections and herbaria as well as robust field sampling. Furthermore, sequencing of COX1 from mycosed scale insect / whitefly cadavers, whether by *Conoideocrella, Dussiella*, or *Aschersonia,* may permit sub-family level identification in the absence of diagnostic morphological features.

The possible occurrence of a paecilomyces-like synanamorph in CK, CT, and DT should also be thoroughly investigated, as should the presence of a hirsutella-like synanamorph in CT. The co-occurence of both types of conidiophores in ARSEF 14590 indicates that both can be produced contemporaneously under certain conditions and might explain the much wider range of conidia measurements for CK, regardless of whether it is a valid *Conoideocrella* sp. or a closely allied novel genus that is yet to be designated.

While both synanamorphs of CL grew sufficiently at ∼20°C, as did the hirsutella-like anamorph of CK, growth rates provided by Mongkolsamrit and colleagues (2016) for CK show significantly reduced growth for the hirsutella-like synanamorph compared to the paecilomyces-like synanamorph for CL reported by Saito and colleagues (2012) and in this study. More work is needed to determine optimal temperature ranges for growth, sporulation, and spore germination for both asexual forms, especially CL’s hirsutella-like synanamorph, which was only recently uncovered. Despite the overlap in growth conditions, each may be adapted for different seasons, host developmental stages, or both given the absence of a sexual stage in the U.S. and Japan. How these temperatures overlap with temperature optima for EHS is also significant. Following the introduction of EHS into the U.S., populations adapted to expand into colder regions (Preisser et al., 2008). While adult female EHS can be found year-round, crawlers are present from spring through autumn. Considering these factors, CL application as a biopesticide may be most effective in early spring, with the aim to establish an epizootic that controls crawlers throughout their emergence.

Though much work has been done on the taxonomy of *Torrubiella* and closely allied fungi over the last decade and a half (Sung et al., 2007; Johnson et al., 2009), some taxonomic uncertainties in and around *Conoideocrella* remain. Most importantly, Petch (1923) noted that he had not examined the type specimen of CL described by Zimmermann (1901) from Java, so how the morphology and phylogenetic placement of this specimen overlaps with our current understanding of CL requires further scrutiny. This is especially relevant since Zimmermann noted that part-ascospores were not observed in Java type material, whereas Petch noted cylindrical part-spores from several locations (Petch, 1923). The placement of *Torrubiella brunnea* also suffers from taxonomic uncertainty (Petch, 1923; Hywel-Jones, 1993).

Despite the urgent need for a major taxonomic revision of *Conoideocrella, Dussiella*, and closely allied genera, it is clear that CL is well-defined as a species and is currently the only *Conoideocrella* species for which pathogenicity has been formally tested. Saito and colleagues (2012) confirmed pathogenicity of CL on *Camellia* spiny whitefly nymphs using three Japanese strains. The results of their bioassays not only revealed significant differences in infection rates across strains and inoculum concentrations, but also that CL outperformed commercially available mycoinsecticides at concentrations below recommended field application rates for those products (Saito et al. 2012). Though pathogenicity was confirmed using our NC CL strains against EHS, reduced infection rates compared to Japanese CL strains on whitefly might be explained by multiple factors: 1) our bioassays were conducted over a 2-week period compared to a three week period for whitefly bioassays, which showed significant jumps in infection rates between week 2 and 3; and 2) our bioassay used a single (medium) concentration treatment that failed to cause higher infection rates against whiteflies using Japanese CL strainsfor two of three strains compared to the negative control after 2 weeks (Saito et al., 2012). Given the close phylogenetic relationship between U.S. and Japanese strains, it is plausible that whiteflies may serve as the preferred host over armored scale insects despite the confirmed susceptibility of various members of Coccidae, Diaspididae, and Aleyrodidae. Given the numerous records from the southeast from whiteflies between 1913-1939, surveys to recover and characterize CL from whiteflies including assessing pathogenicity is urgently needed.

The lack of reports of CL especially from whiteflies from the early 1940s through the 2010s is perplexing. One possible explanation is that CL may have been mistaken for another entomopathogenic fungal genus that infects whiteflies, *Aschersonia,* which has been reported widely in the southeast U.S since the early twentieth century. Both produce orange-colored stroma atop infected insects on the underside of leaves (Fasulo & Weems, 1999; Martini et al., 2022). Otherwise, the absence of CL observations during this prolonged time period may reflect: a gradual, albeit unchecked decades long host expansion of CL from whiteflies in the southeastern U.S. to armored scale insects possibly due to declining citrus whitefly populations; a precipitous drop in EHS’s primary host (eastern hemlock) due to mortality from hemlock woolly adelgid across most of the introduced range of EHS; 3) a relatively recent jump by EHS to Christmas trees (firs) that may support higher densities of EHS or unique microsite conditions compared to eastern hemlock, thus allowing for the emergence of an epizootic; 4) an increase in classical biocontrols including *Encarsia citrina*, a parasitoid wasp of adult EHS, and *Cybocephalus nipponicus, a* beetle predator of EHS, which were found among CL-infected EHS in North Carolina and may spread spores; and/or 5) changes in management, particularly the use of insecticides that impacted fungal growth.

Insect growth regulators like buprofezin have provided the most consistent and long lasting control against EHS in Christmas tree farms (Sidebottom, 2016). Coincidentally, comparisons among 17 commercial insecticides on CL growth showed buprofezin and 11 other insecticides significantly reduced fungal growth (Saito et al. 2012), which might account for the lack of previous observations in these treated stands. Interestingly, most of the sites where CL was detected were on farms where management had been abandoned, including the use of insecticides. On the other hand, imidacloprid, which is commonly used to treat HWA and EHS, had no effect on growth of CL (Saito et al. 2012), assuming adequate healthy populations of EHS exist in an area to sustain fungal populations. The preferential use of imidacloprid in hemlock forests and buprofezin in Christmas tree farms may allow CL to survive in areas adjacent to Christmas tree farms.

Our study provides important historical context for understanding this modern CL epizootic among EHS in U.S. Christmas tree farms. CL has been observed infecting insects across the eastern U.S. since the early 1900s. The observed host range of CL includes multiple economically important pest insects, but lacks insects that would be considered beneficial. The primary spore CL produces in artificial culture is infective to the primary motile stage of EHS, which is key for the spread of this pest insect in Christmas tree farms. Motile stages are also more likely to encounter an applied biopesticide. Multiple factors, including Christmas tree management practices, may contribute to recent and conspicuous outbreaks among U.S. EHS populations. Considering the developmental stage specificity of CL, its integration as a biocontrol intervention against EHS would be most effective just before and throughout crawler emergence. Development of CL for applied use by the Christmas tree industry may help alleviate the burden of EHS on tree export; however, the potential of this product may extend beyond Christmas tree orchards controlling other pest insects.

## Acknowledgments

We acknowledge that this work was completed on the traditional land of the Osage, Shawnee, and Massawomeck peoples. Samples were additionally collected on the land of Cherokee, Yuchi, and Moneton peoples. Special thanks to Ryan Percifield, Director of the West Virginia University Genomics Core Facility, who assisted with library prep and sequencing of *C. luteorostrata* strain ARSEF 14590. This project was funded by the Christmas Tree Promotion Board Grants #20-10-WVU and #21-07-WVU. BL was supported by the USDA (USDA-ARS Project 8062-22410-007-000D). HB was supported by WVU’s Research Apprentice Program, and the honors Biology capstone program, under Dr. Susan Raylman in the West Virginia University Department of Biology. AMM was supported by a WVU Outstanding Merit Fellowship. JES is a CIFAR fellow in the program Fungal Kingdom: Threats and Opportunities and was supported by the National Science Foundation (NSF) (EF-2125066) and the U.S. Department of Agriculture (USDA) (National Institute of Food and Agriculture Hatch projects CA-R-PPA-211-5062-H). Analyses were performed on the UC Riverside High Performance Computing Cluster supported by the NSF (DBI-1429826 & DBI-2215705) and the National Institutes of Health (NIH) (S10-OD016290).

## Supplemental information

Supplemental code and files are available at https://github.com/HanaBarrett/EHS-CL-Analysis.

SUPPLEMENTARY FIG. 1. EF1-α phylogenetic tree for the Clavicipitaceae dataset. Topology and branch lengths shown are from the ML analysis. Bootstrap support and posterior probabilities are indicated for each node supported in the ML analysis (ML/BI). Strain metadata including GenBank accession numbers are listed in Table 2.

SUPPLEMENTARY FIG. 2. 28S phylogenetic tree for the Clavicipitaceae dataset. Topology and branch lengths shown are from the ML analysis. Bootstrap support and posterior probabilities are indicated for each node supported in the ML analysis (ML/BI). Strain metadata including GenBank accession numbers are listed in Table 2.

SUPPLEMENTARY FIG. 3. RPB1 phylogenetic tree for the Clavicipitaceae dataset. Topology and branch lengths shown are from the ML analysis. Bootstrap support and posterior probabilities are indicated for each node supported in the ML analysis (ML/BI). Strain metadata including GenBank accession numbers are listed in Table 2.

SUPPLEMENTARY FIG. 4. RPB2 phylogenetic tree for the Clavicipitaceae dataset. Topology and branch lengths shown are from the ML analysis. Bootstrap support and posterior probabilities are indicated for each node supported in the ML analysis (ML/BI). Strain metadata including GenBank accession numbers are listed in Table 2.

## Notes

https://github.com/HanaBarrett/EHS-CL-Analysis

### Competing Interest Statement

JES was a paid consultant for Zymergen, Sincarne, and Michroma

### Summary of Updates

This revision includes: 1) Additional phylogenetic analyses using additional loci and strains, 2) a comprehensive summary of all previous records of C. luteorostrata, other Conoideocrella spp. and Dussiella spp. around the globe, 3) Addition morphological data and comparisons including the discovery of a hirsutella-like synanamorph in C. luteorostrata.

